# Deep Learning Enabled 3D Multi-Omic Analysis Reveals Molecular Signatures of Heterogeneous Response to Chemotherapy in Pancreatic Cancer

**DOI:** 10.64898/2026.03.03.709150

**Authors:** André Forjaz, Hengameh Mojdeganlou, Alens Valentin, Meredith Wetzel, Dmitrijs Lvovs, Atul Deshpande, Sarah M. Shin, Sujan Piya, Kimal I Rajapakshe, Paola A Guerrero, Brian A Pedro, Dimitrios N Sidiropoulos, Pei-Hsun Wu, Vincent Bernard Pagan, Demystifying Pancreatic Cancer Therapies TeamLab, Denis Wirtz, Elana J Fertig, Luciane T Kagohara, Won Jin Ho, Ashley L Kiemen, Laura D Wood

## Abstract

Resistance to systemic therapy is a major unmet challenge in pancreatic cancer. To identify potential mechanisms of resistance, we developed a novel 3D pipeline in clinical samples that uses deep learning to classify sensitive and persistent tumor cell populations based on morphological features, enabling subsequent molecular characterization of intratumoral heterogeneity. We applied this automated 3D pipeline to a cohort of human pancreatic cancer samples treated with neoadjuvant chemotherapy, identifying heterogeneity in response to therapy both between and within tumors. Application of spatial proteomics to these sensitive and persistent regions identified enhanced epithelial-to-mesenchymal transition and non-classical cell states in persistent cells, confirming our morphological classification. Integration of spatial transcriptomics in multiple pancreatic cancer cohorts associated fibroblast-cancer crosstalk via syndecans with resistance to cytotoxic therapy. Our validated 3D multi-omic pipeline is now poised for application to clinical trials, enabling discovery of resistance mechanisms and design of new therapeutic combinations to circumvent resistance.

**Statement of significance:** We developed a novel 3D multi-omic pipeline to identify mechanisms of resistance to chemotherapy in clinical samples. This approach associated fibroblast-cancer crosstalk via syndecans with resistance to cytotoxic therapy and is poised for broader application in neoadjuvant clinical trials.

## Introduction

Pancreatic ductal adenocarcinoma (PDAC) is a highly lethal cancer that is predicted to become the second leading cause of cancer death in the United States by 2030 (1). Improvements in systemic therapy are critical to lengthening survival in this disease. Accomplishing this goal requires deeper understanding of the molecular and cellular mechanisms that drive therapeutic resistance, to both standard-of-care chemotherapy and to novel therapies currently in clinical trials. Although chemotherapy resistance is pervasive in PDAC, mixed responses to systemic therapy are common, indicating that subsets of cancer cells within the same patient (or even within the same tumor) may have differing sensitivity to therapy. Analysis of human tissue samples after systemic therapy can provide insights into mechanisms of intra-tumoral response and resistance; however, leveraging the full potential of these samples requires the development of new approaches integrating anatomical, cellular, and molecular analysis approaches.

To identify molecular and cellular features associated with response and resistance to chemotherapy in pancreatic cancer, we designed a novel, deep learning-based workflow with integrated multi-omic analyses. This workflow uses quantitative 3D imaging and tumor cell morphology to identify sensitive and persistent cancer cell subsets, followed by spatial analysis of heterogeneous regions of interest to characterize molecular and cellular features of response, resistance, and crosstalk with the surrounding microenvironment. Application of this new pipeline nominates novel potential mechanisms of pancreatic cancer resistance to cytotoxic therapy. In addition, our approach represents a powerful tool for facile integration into clinical trial pipelines to rapidly identify mechanisms of resistance that can inform putative therapeutic combinations for application in subsequent preclinical and clinical studies.

## Results

### A novel deep learning-based 3D workflow enables targeted characterization of treatment response

Characterizing intra-tumoral heterogeneity in therapeutic response often requires intensive manual histopathological assessment of multiple regions of PDAC resections, which can be more effectively completed through 3D characterization and automated annotation. To enable this more automated and comprehensive annotation, we prospectively collected thick blocks of formalin-fixed, paraffin embedded (FFPE) tumor tissue from 13 individuals who underwent pancreatic resection following a diagnosis of PDAC. Two samples were collected from treatment naïve patients, and 11 samples were collected from patients who received neoadjuvant chemotherapy prior to surgery (see clinical details in **Table S1**). Blocks were serially sectioned, and every third slide was stained with hematoxylin and eosin (H&E) and digitized (**Fig. 1A**). We used the CODA software to perform nonlinear image registration to generate semi-continuous stacks from the discrete H&E images (2). We trained a semantic segmentation model to recognize ten components of the pancreatic microanatomy in the histology images: PDAC, non-neoplastic ducts, acini, islets, smooth muscle, endothelium, nerves, immune cell aggregates, stroma, fat, and background (**Fig. 1B**). In independent testing, the model achieved 96.6% overall accuracy (**Fig. S1A**). To incorporate nuclear morphology, we used StarDist to extract 20 morphological features (including area, major axis length, and color) per nucleus (**Fig. S1B**) (3,4). This workflow enabled us to generate 3D cellular-resolution tumor maps by integrating the registration transforms with the tissue segmentation masks and nuclear features (**Fig. S1C**).

**Figure 1.**
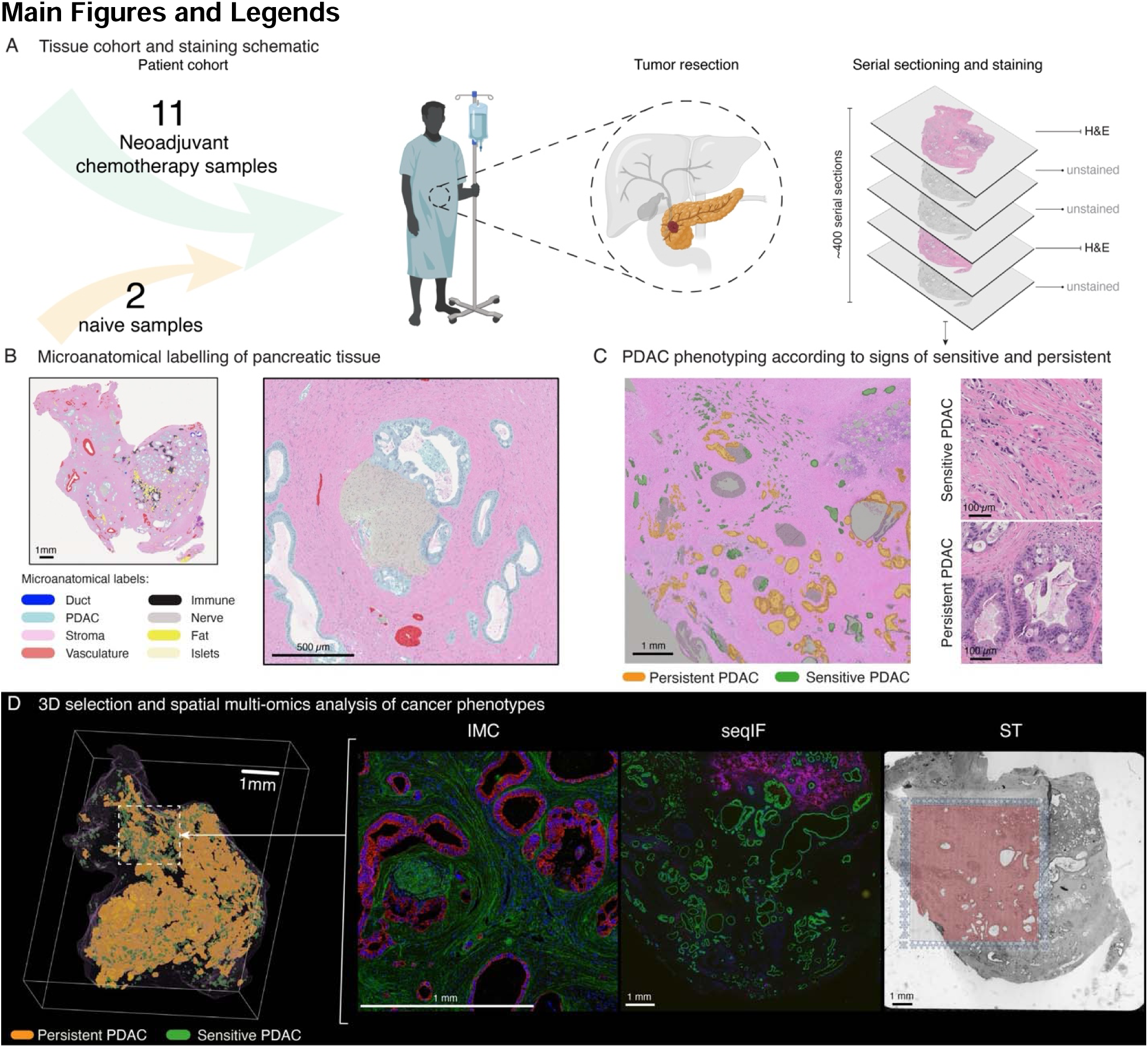
A novel workflow for targeted characterization of treatment response in pancreatic cancer. **(a)** Treated and untreated tumor samples collected from pancreatic cancer patients were serially histologically sectioned for 3D reconstruction. **(b)** A deep learning model was trained to detect ten pancreatic microanatomical components. **(c)** Sensitive and persistent tumor cells were defined using 3D connectivity and correspond to 2D cellular morphology. **(d)** The 3D distribution of sensitive and persistent cancer populations was used to guide placement of spatial proteomics and transcriptomics for targeted molecular profiling of heterogeneous treatment response regions.

Conventional two-dimensional (2D) diagnostic grading of treatment response to cytotoxic therapy in PDAC relies on the size and connectivity of tumor cell groups, with single cells or small clusters indicative of response to therapy (5). Although this approach has limitations related to interobserver variability, these 2D morphological features have been associated with outcome after neoadjuvant chemotherapy, with single cells or small clusters correlated with improved prognosis (6). These data suggest that cancer cells that are sensitive to therapy can be distinguished from unaffected, or persistent, cancer cells through assessment of their histopathologic features. Specifically, sensitive cancer cells appear as disconnected cell clusters with more abnormal cellular features compared to persistent cancer cells, which appear as larger, interconnected structures (**Fig. 1C**). We hypothesized that these observations translate to 3D, and that we could reliably distinguish sensitive and persistent cancer cells using cancer cluster volume. We used 0.01 mm^3^ as a volume cutoff, which differentiated large connected glandular objects and discrete cell clusters upon visual inspection. In addition to comparing the composition of sensitive and persistent cancer cells across our cohort, we also aimed to use their spatial distribution in 3D to guide selection of regions of perceived heterogeneous response for downstream molecular analysis (**Fig. 1D**).

In 3D, the cancer populations appeared intermixed, with large, connected regions of persistent cancer clusters interspersed with sensitive cancer clusters (**Fig. 2A**). To validate that our volume-based definition of sensitive and persistent cancer clusters captured distinct tumor cell populations, we compared their nuclear morphological features. This revealed that sensitive cancer cell nuclei had significantly larger area, longer perimeter, and more heterogeneous hematoxylin staining compared to persistent cancer cell nuclei (**Fig. 2B**). Importantly, these differences in nuclear morphology are in line with pathologist observations of cellular changes in chemotherapy-treated samples (7,8). We observed notable interpatient heterogeneity, with some tumors primarily containing persistent cells and others containing large proportions of both sensitive and persistent cells. As expected, untreated samples contained predominantly persistent cancer cells, supporting our hypothesis that the cell clusters labeled as sensitive were a feature mainly present in chemotherapy-treated tissue. These observations were supported by quantitative analysis, as total sensitive and persistent sample volume and composition show high heterogeneity in treated samples, and very low sensitive tumor presence in treatment naïve samples (**Fig. 2C,D**).

**Figure 2.**
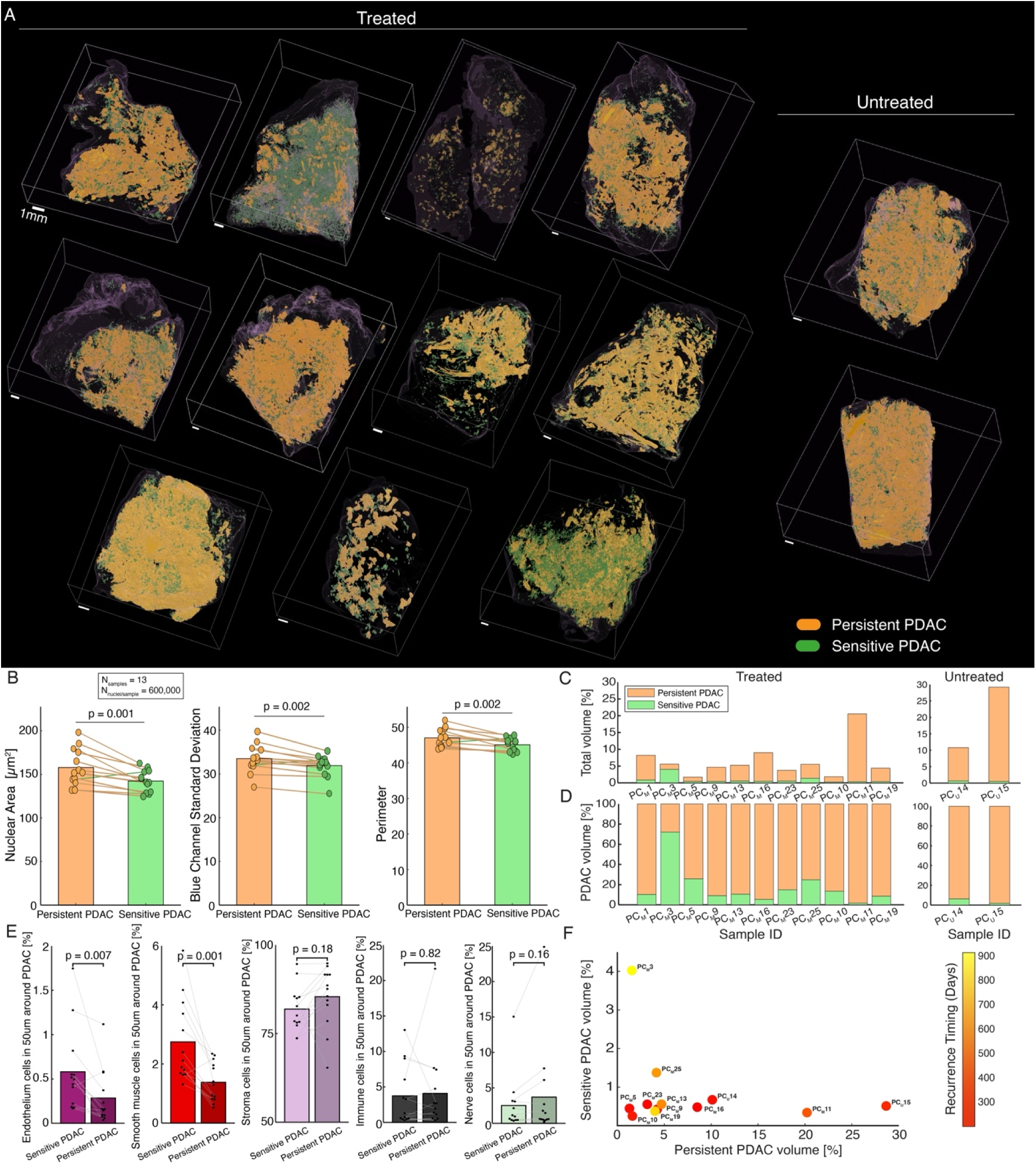
**Quantitative evaluation of sensitive and persistent tumor cells**. **(a)** 3D renderings of sensitive and persistent pancreatic ductal adenocarcinoma (PDAC) in 13 human tumor samples display complex spatial organization and high inter-patient variability. **(b)** Bar plots comparing the mean per-sample nuclear area (p=0.001), standard deviation in blue-color intensity (p=0.002), and nuclear perimeter (p=0.002) between sensitive and persistent tumor cells across the thirteen 3D samples. Significance calculated using a paired t-test (with paired points from the same tumor connected by lines in the bar plots). **(c-d)** Bar plots depicting the percent composition of sensitive and persistent cancer normalized by (c) total sample volume and (d) total cancer volume. **(e)** Bar plots comparing the mean per-sample composition of cells in the 50 micron surrounding sensitive and persistent cancer cells. Shown are endothelium (p=0.007), smooth muscle (p=0.001), stroma (ns), immune hotspots (ns), and nerves (ns). Significance calculated using a paired t-test. **(f)** Scatter plot depicting the per-sample percent composition of sensitive cancer as a function of percent composition of persistent cancer. Marker color represents the corresponding time to recurrence following surgical resection. Note: Sample PC_M_1 is excluded from (f) as the patient died 90 days post-surgery without recurring. Sample PC_M_19 is labelled at recurrence time 770 days as this is the latest follow-up time. However, the patient has not yet recurred.

Next, we compared the cellular neighborhoods surrounding the sensitive and persistent cells. While the composition of stromal cells, nerves, fat, exocrine, and endocrine cells was similar, there was a higher composition of endothelial and smooth muscle cells within a 50 µm radius surrounding the sensitive cancer cells, suggesting sensitive cancer cells were more closely associated with vasculature compared to persistent cancer cells (**Fig. 2E**). Notably, this finding is consistent with previous literature suggesting that delivery via the vascular system in a key determinant of response to chemotherapy (9,10).

To determine the impact of cancer cell composition on patient outcome, we next assessed the persistent and sensitive tumor cell volume in the context of time to post-operative tumor recurrence (**Fig. 2F**). We found that the two samples containing the highest percentage of persistent tumor cells came from patients who experienced rapid recurrence (within 1 year), while the sample containing the highest percentage of sensitive cancer cells came from the patient with the longest time to recurrence (900 days). However, there was no statistically significant correlation between time to recurrence and composition of sensitive and persistent cancer cells when considering the entire patient cohort. Taken together, these results suggest that while there were striking associations in individual tumors, cancer cell composition alone did not predict clinical outcome, at least in our cohort of eleven patients.

After quantifying the proportions of sensitive and persistent cancer cell populations across our cohort, we used these structures to identify regions of interest (ROIs) hypothesized to be highly heterogeneous in their response to treatment. The goal of this analysis was to identify biologically informative regions for in depth spatial profiling to characterize the distinct molecular features of these cancer cell populations and their surrounding microenvironment. Visual inspection revealed that most whole slide images did not contain acceptable regions of mixed cancer cell populations, supporting automated approach to identify ROIs deep within the tumor blocks (**Fig. 3A**).

**Figure 3.**
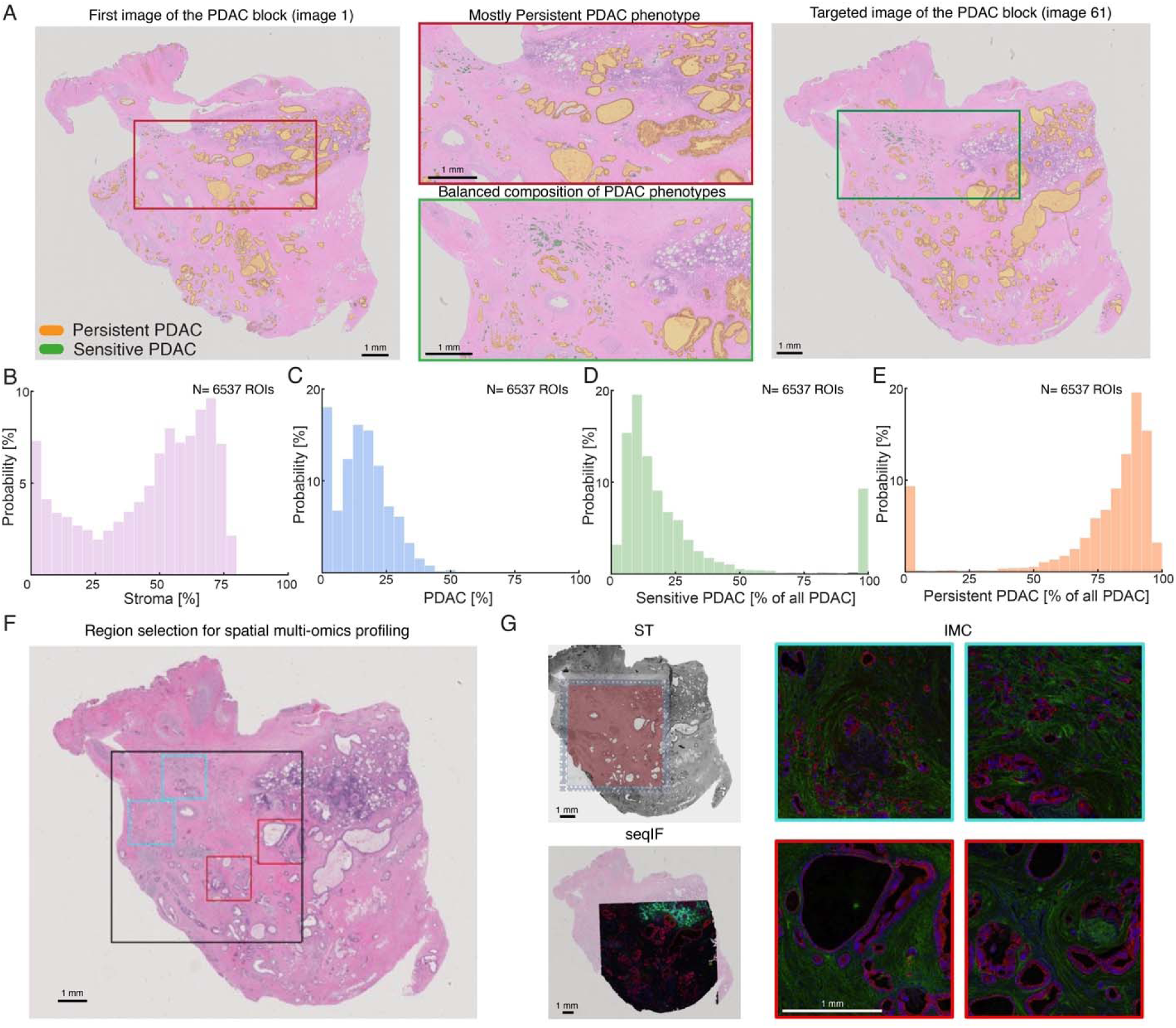
**Automated selection of regions of interest with heterogeneous response**. **(a)** A histology image of the first section of the block with little visible sensitive cancer cells, compared to a histology image taken deeper within the block with a visible mix of sensitive and persistent cancer cells. **(b-e)** Histograms of tissue composition within each of 6,397 regions of interest considered in sample PC_M_1 depicting per-region distribution, or percent probability of (b) stroma composition, (c) cancer composition (d) percent of all cancer that is sensitive, and (e) percent of all cancer that is persistent. **(f)** Example region of interest chosen in sample PC_M_1. The Black square represents the region selected for spatial transcriptomics (ST) and sequential immunofluorescence (seqIF; COMET) profiling, red squares represent regions selected for imaging mass cytometry (IMC) profiling of persistent cancer cells, and blue squares represent regions selected for IMC profiling of sensitive cancer cells. **(g)** Example images taken from the ST, seqIF, and IMC platforms correspond to the target regions outlined in (f).

To provide comprehensive analysis of both cell types and cell-cell crosstalk in the identified high-information regions, we chose to leverage two complementary spatial proteomics technologies, imaging mass cytometry (IMC) and sequential immunofluorescence (seqIF), as well as spatial transcriptomics (ST) on the Visium platform. Importantly, each of these assays had distinct parameters to define analyzable regions. To select regions to analyze with all three approaches, we first simulated extraction of 6.5 x 6.5 mm^2^ capture window areas across all whole slide images in each 3D reconstructed sample. Within each capture window, we calculated the tumor percentage and ratio of sensitive and persistent tumor cells to identify an ROI containing abundant numbers of each (**Fig. 3B-E**). As ST and seqIF allowed for imaging of one large capture window, we targeted a region containing both sensitive and persistent cancer cells. As IMC allowed capture of four small 1.5 x 1.5 mm^2^ windows within the original 6.5 x 6.5 mm^2^ ROI, we next targeted two ROIs towards regions enriched for persistent cancer cells and two ROIs towards regions enriched for sensitive cancer cells (**Fig. 3F,G**). Altogether, this method allowed us to identify images containing mixed sensitive and persistent cell populations for deeper profiling.

### High-parameter spatial proteomics defines tumor cell and microenvironmental features of persistence after chemotherapy

To deeply characterize sensitive and persistent tumor cells with their surrounding microenvironment, we performed high-parameter spatial proteomics on CODA-defined ROIs containing both cancer cell populations. Specifically, IMC was performed using a 45-channel panel (**Fig. 4A**) designed to resolve major cellular compartments, functional immune states, and tissue architecture within the tumor microenvironment (TME) by high-fidelity, protein-level phenotyping. The panel included immune lineage (CD45) and functional markers (GZMB, PD-1, PD-L1, IL-6, IL-8, CXCL12) to define lymphoid (CD3, CD4, CD8, CD20, CD45RA, CD45RO, FOXP3, CD57) and myeloid (CD68, CD163, DC-SIGN, CD16) populations and their activation states; antigen-presenting cell markers (HLA-DR, CD74) to identify APC subsets; and proliferative marker Ki-67. Stromal/mesenchymal markers (αSMA, Vimentin, Podoplanin, PDGFRα, FAPα, Collagen, S100A4, N-cadherin, CD105) and vascular markers (CD31) were used to characterize fibroblast and endothelial compartments, respectively. Epithelial and differentiation markers (pan-cytokeratin, E-cadherin, KRT17, S100A2, TFF1, GATA6) enabled identification of cancer cell states. Architectural and segmentation markers (DNA intercalators, membrane and cytoplasmic markers) supported accurate cell segmentation and spatial analysis. Acquired images displayed major cell types as intended (**Fig. 4B**). Image segmentation from 16 total regions yielded 125,091 cells, which were then clustered into a total of 25 annotated clusters (6 cancer, 9 lymphoid, 6 myeloid, 3 fibroblast, and 1 endothelial; **Fig. 4C**), enabling a more detailed phenotyping of cells, complementing what could be achieved by CODA. Specifically, cancer cells were subtyped based on protein markers selected to annotate previously established states (11), including “classical” (positive for TFF1 and/or GATA6), “basal” (positive for KRT17 and/or S100A2), “mixed” (any mixture of classical and basal markers), and cancer cells not otherwise specified, “cancer_NOS”, due to limited expression of subtyping markers. IMC also delineated CD4^+^ helper (Th), FOXP3^+^ regulatory (Treg), GZMB^+^ CD45RA^+^ cytotoxic effector (Tc_Eff), and CD45RO^+^ memory subtypes of T cells; CD163^lo^ (Mac_I) and CD163^hi^ (Mac_II) macrophage subtypes; and cancer-associated fibroblast (CAF) subtypes including collagen^hi^ CD105^hi^ (CAF_I), collagen^hi^ LSMA^hi^ (CAF_II), and CXCL12^hi^ (CAF_III). As expected, CAF and myeloid subtypes were consistently the most abundant components, making up approximately 40% and 30% on average, respectively, within the PDAC TME. Within the cancer cell compartment, proportional abundances skewed toward non-classical subtypes (average of 62% across all four tumors). While we observed both intra- and inter-tumoral heterogeneity across all regions analyzed, cancer cell and TME compositions from multiple regions within the same tumor were notably more similar (**Fig. 4D**).

**Figure 4.**
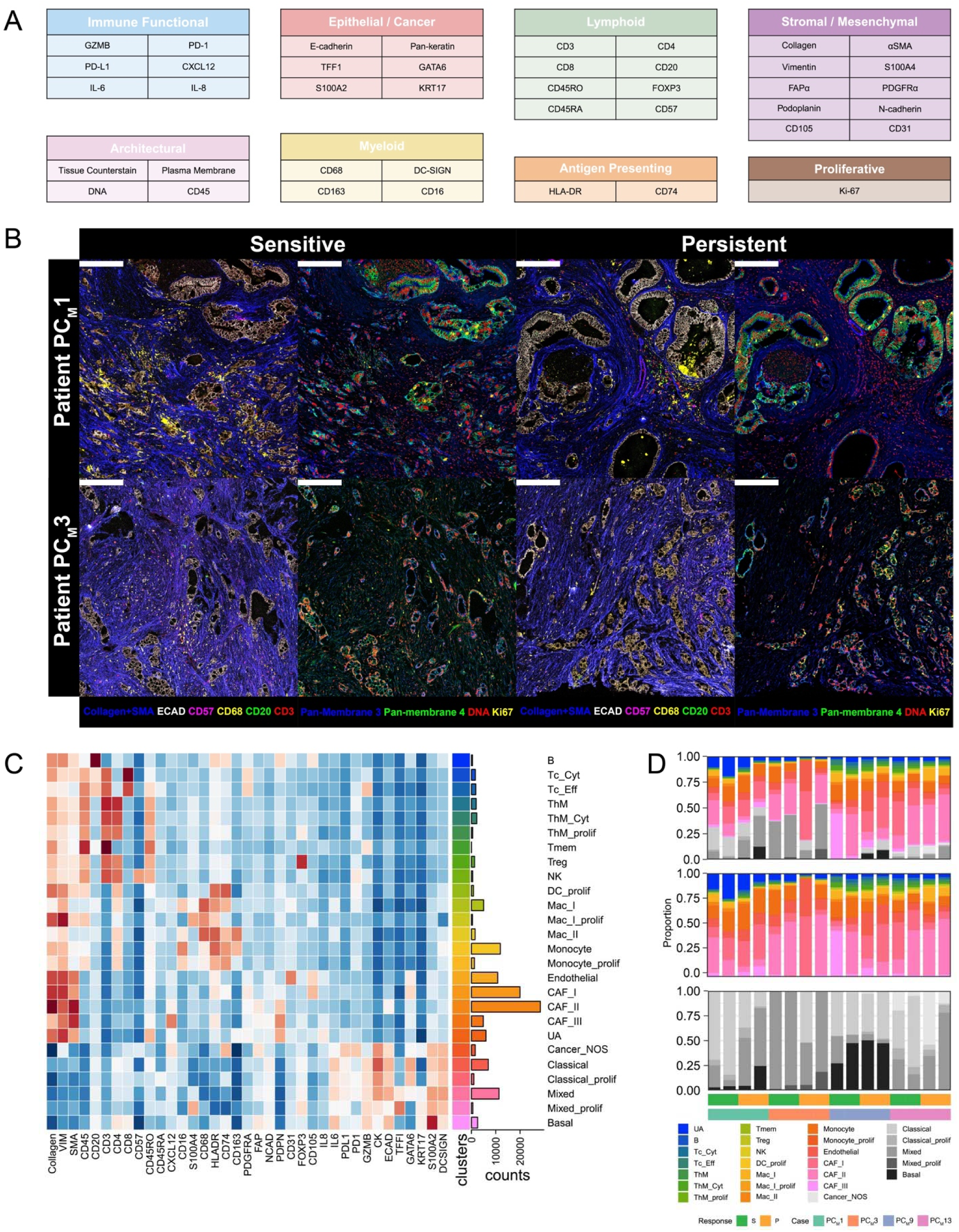
IMC analysis yields detailed cellular subtypes within CODA-selected regions with sensitive and persistent PDAC. **(a)** IMC panel of 45 channels incorporating 41 antibodies (plus other architectural reagents such as tissue counterstain and DNA intercalator) optimized for characterizing epithelial / cancer cells, stromal / mesenchymal cells, lymphoid cells, myeloid cells, and functional states. **(b)** Representative images from two unique patients demonstrate quality staining results for tumor regions that are sensitive to treatment and those that persist post-therapy. Scale bar: 300 µm. **(c)** Segmented single-cell dataset was clustered using PhenoGraph and merged into 26 annotated cell types. Heatmap shows a scaled expression profile. **(d)** Stacked bar plots display cellular compositions of all (top), non-epithelial (middle), and cancer (bottom) subtypes per region analyzed. Abbreviations: Tc, cytotoxic CD8^+^ T cells; Th, helper CD4^+^ T cells; Tc_Cyt, GZMB^int^; Tc_Eff, CD45RA^+^GZMB^hi^; ThM, CD4^+^CD45RO^+^ memory T cells; Tmem, CD3^+^CD4^-^CD8^-^ memory T cells; Treg, CD4^+^FOXP3^+^ regulatory T cells; NK, CD57^+^ natural killer; DC, dendritic cells; Mac, macrophages; CAF, cancer-associated fibroblasts; prolif, Ki67^+^ proliferating; NOS, not otherwise specified; UA, unassigned.

We further validated these detailed observations using an orthogonal single-cell proteomic approach based on seqIF (COMET). The 20-marker panel used for seqIF was designed to have an overlap of key markers for canonical phenotyping. However, as an independent verification of antibody-based high-parameter profiling, alternative yet well-published antibody clones were selected when readily feasible (pan-cytokeratin, TFF1, KRT17, CD3, FOXP3, CD163), and different markers altogether were included to enhance cell type capture, e.g., CD11c for dendritic cells and CD66b for neutrophils (**Fig. S2A**). seqIF-based single-cell profiling largely recapitulated the results obtained through IMC, highlighting the predominance of CAF and myeloid cell types as well as inter-tumor heterogeneity (**Fig. S2B**). Despite the use of different antibody clones for TFF1 and KRT17, cancer cell subtype compositional profiles were also largely consistent across tumors. An exception was observed in PC_M_1, where omission of the basal subtype marker S100A2 from the seqIF panel resulted in limited detection of mixed phenotypes (**Fig. S2C**,**D**). Overall, these findings established a high level of rigor in phenotypes delineated by our proteomic analysis.

Previous studies have implicated epithelial-to-mesenchymal transition (EMT) signatures, including high vimentin expression, with resistance to chemotherapy in PDAC (12–14). We explored whether our phenotype-rich IMC dataset, which included quantitative vimentin assessment at a per-cell level (**Fig. 5A**), could recapitulate these prior observations when comparing CODA-determined regions of sensitivity and persistence to chemotherapy. Indeed, regardless of the cancer subtype annotation, significantly higher vimentin expression was noted in persistent cancer cells (**Fig. 5B, Fig. S3**), suggesting that our histology-based classification identified molecularly distinct subtypes of tumor cells. More recently, COMPASS trial suggested that the basal PDAC subtype is more resistant to chemotherapy (15,16). In our region-based analysis, we also found that persistent regions harbored cancer cells with higher expression of basal subtype markers KRT17 and S100A2 (**Fig. 5C**). We detected significantly higher density (i.e., number of cells per tissue area) of total cancer cells and trends toward higher densities of non-classical subtypes in persistent regions (**Fig. 5D**). Since binary designation of regions as “sensitive” or “persistent” limited the analytical resolution of chemotherapy responses, we also leveraged CODA quantification to generate a continuous metric of persistence after chemotherapy (i.e., % of persistent cancer cells in a region). This higher resolution analysis demonstrated that the density of mixed phenotype cancer cells significantly correlated with persistence after chemotherapy (R=0.73, p=0.0012, **Fig. 5E**). These results corroborated the presence of non-classical cancer cells as an important axis of chemotherapy resistance, illustrating the power of deep learning-based morphological analysis to uncover clinically significant molecular subtypes.

**Figure 5.**
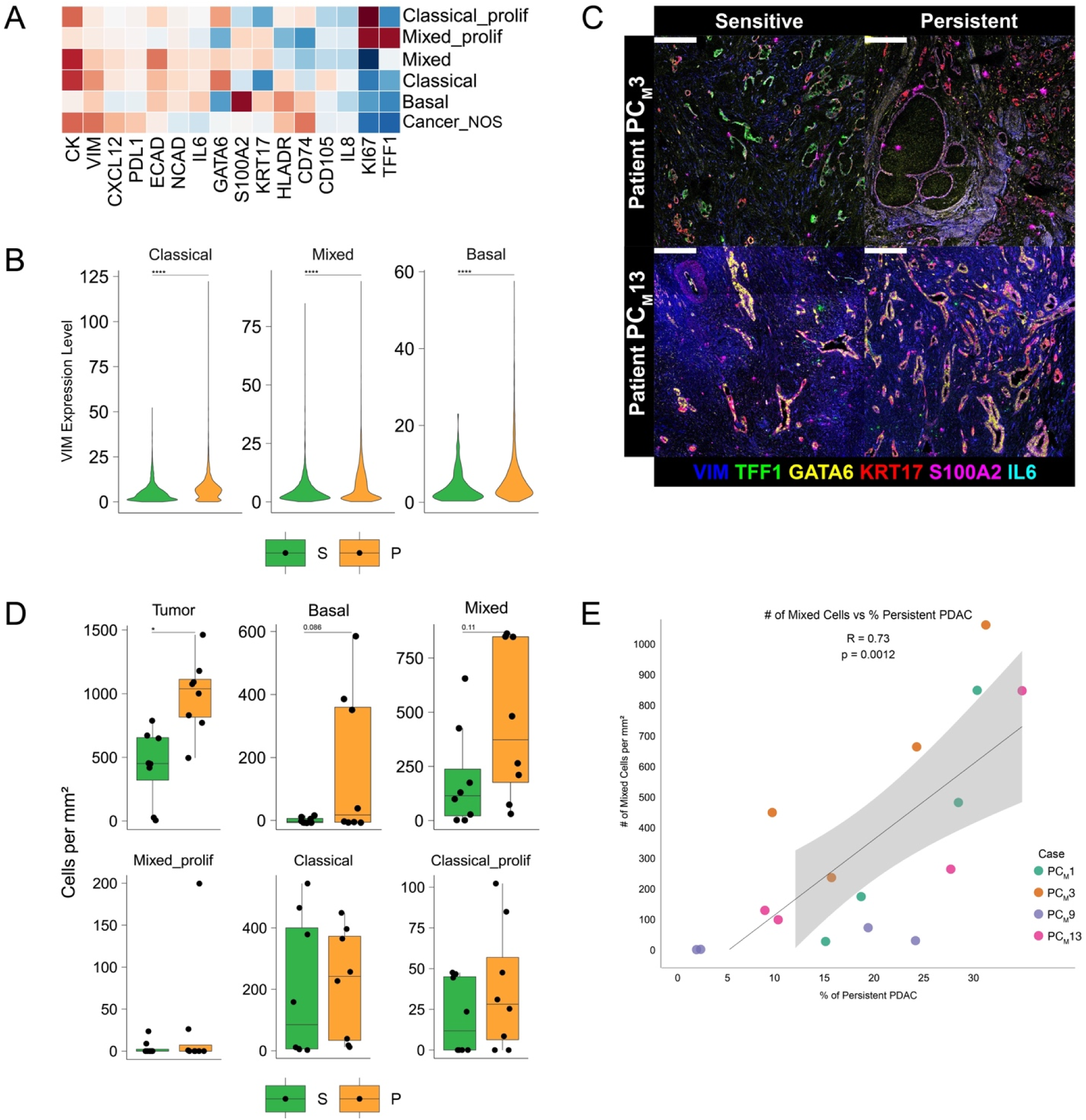
Deep protein-level phenotyping analysis with IMC recapitulates signatures of chemotherapy resistance. **(a)** A heatmap of cancer subtypes with scaled expression profiles. **(b)** Violin plots of per-cell expression of vimentin, a marker of EMT, by classical, mixed, and basal cancer subtypes in CODA-determined sensitive (S) vs. persistent (P) regions compared by unpaired t-test. ****p<0.0005. **(c)** Representative IMC images of cancer subtyping markers. Scale bar: 300 µm. **(d)** Boxplots showing density of cancer subtypes compared by unpaired t-test. *p<0.05. **(e)** Dotplot displaying correlation between CODA-derived calculations of persistent cancer cells and density of mixed cells within each region. P-value calculated by Pearson correlation.

To nominate candidate microenvironmental regulators of cancer chemotherapy responses, we first quantified the spatial influence of neighboring cell types around cancer cells using FunCN (17), which accounted for both the abundance and proximity of each cell type in the cellular neighborhood. Among all cell types, CAFs by far had the highest spatial influence around cancer cells (**Fig. S4**), warranting a deeper evaluation of CAF-PDAC interactions, which we performed using spatial transcriptomics.

### Spatial transcriptomics identifies CAF signaling through syndecans associated with EMT activation in PDAC cells that are non-responsive to chemotherapy

The imaging and proteomics analyses of our 3D-mapped cohort demonstrated that the co-localization of CAFs was associated with persistent cancer cells after chemotherapy. While powerful for inferring cellular co-localization, the spatial proteomics assays were limited in unbiased discovery to enable inference of downstream activation of signaling pathways resulting from cell-to-cell communication. In addition, the heterogeneous response within each 3D tumor sample limited our ability to directly link the cellular networks to patient-level therapeutic response. To provide a complementary dataset, we sought to first assess cell-cell crosstalk in a new cohort of PDAC samples selected for homogeneous histopathological response and well-annotated clinical outcome after neoadjuvant chemotherapy. While such a homogeneous response is not typical in PDAC, this cohort provided a powerful tool in which to identify crosstalk mechanisms associated with clinical response, which could then be further interrogated in our 3D-defined sensitive and persistent cell populations. This new cohort consisted of six PDAC samples with homogeneous histopathological response and concordant clinical outcome, three “responding” PDACs (with pathological response and no recurrence) and three “non-responding” PDACs (with no pathological response and rapid recurrence) (see clinical details **Table S1**). Thus, rather than comparing within patients as in the previous 3D analysis, this new cohort allowed us to compare between patients to identify CAF-PDAC crosstalk mechanisms that mediated distinct responses to chemotherapy and clinical outcome. In addition, consistency between the two distinct cohorts could further confirm our anatomical classification of sensitive and persistent cancer cells by demonstrating similarities in features associated with clinical outcome.

To isolate the intercellular interactions between CAFs and PDAC cells associated with clinical response to chemotherapy, we performed Visium High-Definition (HD) spatial transcriptomics (10x Genomics) on the new PDAC cohort described above. To analyze the resulting data, following sequencing and data preprocessing, we applied Robust Cell Type Decomposition (RCTD) (18) to the Visium HD 16mm bins to annotate detected cell types using a PDAC single-cell RNA-sequencing atlas as a reference (19) (**Fig. 6A, B** and **Fig. S5**). This approach also enabled annotation and removal of acellular bins containing primarily extracellular matrix deposition. To identify signaling pathways associated with lack of response to chemotherapy in PDAC, we performed differential expression and gene set enrichment analysis for the spatial bins annotated as PDAC, specifically comparing the PDAC cells from the “responding” samples to those in the “non-responding” samples. This differential expression analysis demonstrated that across previously defined PDAC transcriptomic subtypes (20) (classical, intermediate, basal; **Fig. S6**) the epithelial-mesenchymal transition (EMT) pathway was enriched in non-responders, independent of the transcriptional PDAC subtype classification (**Fig. 6C**). Hence, while the basal subtype has been commonly associated with enhanced resistance to chemotherapy and increased expression of EMT marker genes, our observation suggested subtype-independent chemotherapy resistance associated with mesenchymal-like features even in the classical subtype.

**Figure 6.**
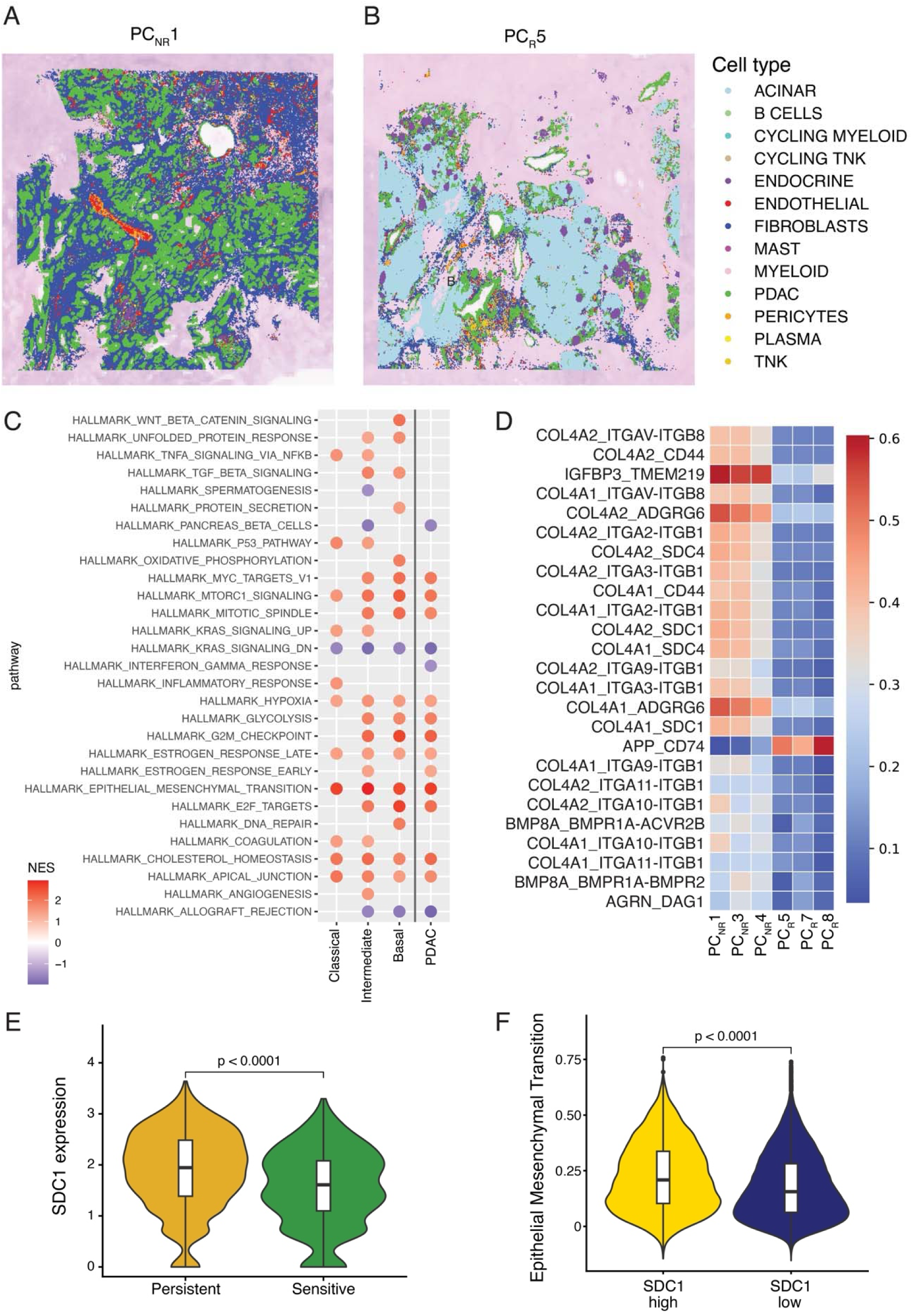
**Spatial transcriptomics analysis of chemotherapy response in PDAC**. **(a-b)** Representation of cell type classifications from Visium HD data in a non-responding (PC_NR_1) and responding sample (PC_R_5). **(c)** Pathway enrichment analysis across PDAC subtypes (classical, intermediate, basal) and between “non-responding” and “responding” PDAC. **(d)** Ligand-receptor pairs identified from the intercellular interaction analysis of CAF to PDAC as differentially activated in “non-responding” compared to “responding” PDAC. **(e)** SDC1 expression in persistent and sensitive PDAC spots with mixed detection of PDAC and CAF from 3D-mapped cohort. **(f)** EMT signaling pathway expression relative to SDC1 high and SDC1 low expression on CAF and PDAC mixed spots from 3D-mapped cohort.

The results of the spatial proteomics analysis in our 3D-mapped cohort led us to further evaluate the CAF population in the Visium HD data. Specifically, we sought to nominate signaling axes from CAFs to PDAC associated with the lack of response to chemotherapy. Previously, we developed a novel algorithm called SpaceMarkers that compared expression changes when cells were co-localized relative to when they were not to estimate cell-cell communication. SpaceMarkers was initially developed to infer downstream signaling from intercellular communication in Visium standard (SD) data leveraging the co-localization of cells in a single spot to infer communication changes (21). For the current analysis, we adapted SpaceMarkers and the spot-based spatial transcriptomics pipeline to account for the near single-cell resolution of Visium HD. Whereas the previous implementation for SD data relied on spot co-localization, the resolution of Visium HD allowed for inference of expression at a near single-cell resolution. This enhanced resolution provided the potential for the extended SpaceMarkers algorithm to infer the precise cell impacted by the signaling change from cellular co-localization. Thus, we refined the algorithm for directionality of signaling between cells, including notably coordinated signaling from ligands in a sender cell to receptors in a receiver cell that occurred uniquely when these cells were co-localized. We applied the SpaceMarkers approach for Visium HD analysis to infer the ligand-receptor pairs associated with CAF-PDAC interactions in each sample. This provided a score for each pair for statistical comparison between response conditions, allowing us to rank ligand-receptor pairs associated with “responding” and “non-responding” PDAC samples. Comparison of ligand-receptor pairs from the SpaceMarkers analysis of CAF ligands signaling to PDAC receptors identified 25 pairs with differential signaling in “non-responding” PDAC relative to “responding” PDAC (24 pairs upregulated in non-responding PDAC and 1 pair upregulated in responding PDAC) (**Fig. 6D**). Several ligands can signal to multiple receptors, and this analysis identified unique receptors in PDAC cells.

Notably, among the receptors highly expressed in PDAC predicated to interact with CAF ligands, we identified genes from the syndecan (SDC) family, SDC1 and SDC4. Given our observation of EMT in “non-responding” PDAC and previous associations of syndecans with EMT (22), we next investigated whether CAF signaling to PDAC through SDCs was enhanced in the persistent cancer cell regions of our 3D-mapped cohort with heterogeneous response to chemotherapy. Because we already mapped the cellular populations extensively with spatial proteomics assays, we now sought only to isolate the transcriptional signatures in the sensitive and persistent tumor cells and their surrounding CAFs. Therefore, lower-resolution spatial transcriptomics with Visium SD was sufficient for this analysis goal. Cell types captured by the Visium SD spots were annotated using CODA as part of the 3D reconstruction step to classify the sensitive and persistent spots prior to transcriptomic analysis (**Fig. S7**). Subsequently, we applied SpaceMarkers to the 3D-mapped cohort samples, identifying 8 PDAC receptors that were concordant among the SD and HD data, including SDC1 and SDC4, with exact matches for two full ligand-receptor pairs. In the Visium SD data, we observed that the spots containing both CAF and PDAC cells from persistent PDAC regions had increased SDC1 (**Fig. 6E**) and SDC4 (**Fig. S8A**) expression relative to sensitive PDAC regions. Moreover, EMT pathway up-regulation was more frequent in PDAC spatial spots with high SDC1 **(Fig. 6F)** and SDC4 (**Fig. S8B**) expression, supporting that CAF-to-PDAC modulation through this pathway was associated with EMT activation in cancer cells. Altogether, the spatial transcriptomics analyses of two independent cohorts with distinct patterns of chemotherapy response suggested that resistance to chemotherapy in PDAC is potentially driven by CAF-to-PDAC signaling through the SDC family of receptors.

## Discussion

In this study, we report a novel 3D multi-omic tissue processing pipeline that allows identification and deep spatial analysis of persistent and sensitive cancer cell populations after treatment with chemotherapy. By applying this pipeline to tissue samples from surgical resections of PDAC after treatment with standard-of-care neoadjuvant cytotoxic therapy, we identified CAF-tumor crosstalk via syndecans as a molecular mechanism of chemotherapy resistance in PDAC. More broadly, this pipeline represents a valuable tool to be applied in future clinical trials to rapidly identify mechanisms of resistance to novel therapies and iteratively improve combination treatment regimens.

Analysis of human tissue samples from clinical trials conducted in patients with cancer can suggest molecular and cellular mechanisms of therapeutic resistance. While correlative analyses of human tissue samples are commonly performed in many trials, our pipeline provides a unique opportunity to leverage intratumoral heterogeneity to define resistance mechanisms. Multiple previous studies have analyzed human PDAC samples treated with neoadjuvant chemotherapy, but the comparisons in these studies focused on treatment status or clinical outcome (23–25). By defining sensitive and persistent cell populations using quantitative 3D analysis, our pipeline enables intratumoral comparisons to identify molecular drivers of resistance. We leverage intratumoral heterogeneity in therapeutic response that can only be defined by 3D analysis, allowing rapid identification of candidate resistance mechanisms using 3D-guided multi-omic profiling. Importantly, multiple lines of evidence support our anatomical classification, including enhanced epithelial-to-mesenchymal transition (EMT) and mixed basal-classical tumor cells in anatomically defined persistent tumor cells, which have been associated with therapeutic resistance and poor prognosis in previous studies (24,25). Although applied here in the setting of standard-of-care chemotherapy, this pipeline represents an important resource for assessment of clinical trials in the future. By defining potential resistance mechanisms in a relatively small number of samples, this pipeline allows rapid development of new combination therapies that can be tested in subsequent clinical trials. Importantly, our pipeline is also nimble - although the current study leveraged spatial proteomics and transcriptomics, additional assays can be incorporated on intervening sections in our 3D models as guided by the details of the clinical strategy being tested.

The results of our 3D multi-omic analysis suggest that CAF-tumor signaling via syndecans may underpin resistance to cytotoxic therapy. Syndecans are transmembrane proteoglycans with multiple roles in cell-cell and cell-matrix communication. SDC1 has been previously identified to play an important role in pancreatic tumorigenesis, including as a mediator of macropinocytosis upregulated by mutant KRAS and as a mediator of resistance to KRAS-targeted therapy (26,27). Intriguingly, recent work identified SDC1 as a receptor on PDAC cells capable of receiving signals from CAF-derived ligands (28). This study demonstrated an association of high SDC1 expression with poor patient prognosis and showed that SDC1 targeting with an antibody-drug conjugate reduced tumor cell survival and caused extracellular matrix remodeling. Another recent study described targeting SDC1 with a therapeutic antibody, which resulted in pancreatic cancer cell death *in vitro* and enhanced efficacy of cytotoxic therapy against pancreatic tumors *in vivo* in mouse models (29). Targeting SDC1 with lipid-based delivery of siRNA can also inhibit growth of PDAC cells (30), and additional studies have also reported the association of syndecan expression with poor prognosis in pancreatic cancer (31,32). Our study is the first to identify SDC1 as a mediator of resistance to chemotherapy and underscores the potential utility of SDC1 targeting in combination with cytotoxic therapy as well as KRAS inhibition. Multiple approaches to target SDC1 are in clinical development, highlighting the near-term applicability of this strategy (33).

There are multiple limitations to the current study. Due to the cost and throughput of the 3D modeling and comprehensive omics assays, the analyzed cohorts were by necessity small and single center in origin. However, with the available cohorts, we validated the accuracy of our 3D pipeline to distinguish sensitive and persistent cancer cells and identified a mechanism of CAF-PDAC crosstalk associated with resistance to chemotherapy that was correlated with patient outcome in independent cohorts. Second, the Visium leveraged in our 3D-mapped cohort did not have the quasi-single-cell resolution of the HD technology leveraged in our second cohort. Nonetheless, future work could readily integrate this HD technology or other imaging spatial transcriptomics technologies into the pipeline, as we have done previously in the treatment naive context (34–38). However, given the consistency of the findings between the two technologies, we do not anticipate that additional profiling is necessary to delineate pathways underlying the CAF-PDAC interactions in chemotherapy resistance in this case. Future studies should evaluate the relative information gained from distinct technologies for inferring the mechanisms of resistance in our 3D multi-omic integration to minimize cost and assays required for relevant assessment of resistance mechanisms.

This study reports an innovative pipeline for 3D multi-omic tissue analysis to identify mechanisms of therapeutic resistance. Notably, we demonstrated the unique power of deep learning enabled 3D modeling to define sensitive and persistent cancer cell populations associated with intratumoral heterogeneity in therapeutic response. This pipeline is now poised for application, initially in neoadjuvant clinical trials and in the future in later-stage disease platforms, to enable data-driven inference of novel combination therapies to circumvent resistance.

## Materials and Methods

### Acquisition of tissue from surgically resected human pancreatic cancer specimens

For the cohort analyzed with 3D multi-omic profiling, thick (∼1cm) tissue slabs were prospectively obtained from surgically resected PDACs from patients at the Johns Hopkins Hospital treated with or without neoadjuvant therapy. Thirteen total samples were included in this cohort - eleven from patients who were treated with neoadjuvant chemotherapy prior to resection and two from patients who were treatment-naïve at surgery (see clinical details in Table S1). Tissues were formalin-fixed, paraffin-embedded (FFPE), and sectioned to generate one hematoxylin and eosin (H&E)–stained slide for initial histopathologic evaluation by pathologists. This assessment was performed to confirm the diagnosis of PDAC and to verify the presence and adequacy of tumor tissue within each specimen. Following this confirmation, the FFPE blocks were serially sectioned onto several hundred consecutive sections (5 µm thick) ranging from 183-820 slides. Every third section was stained with H&E, while the intervening sections were mounted as unstained plus slides, subsequently stored at -20C for future spatial profiling. H&E-stained slides were scanned and registered using a whole-slide scanner (Hamamatsu nanozoomer S360 and S210) at 20x pixel resolution (∼0.5 µm per pixel) and subsequently utilized for 3D reconstruction using CODA (see below).

For the cohort analyzed by Visium HD alone, all PDAC cases that underwent neoadjuvant therapy were retrospectively reviewed. Six cases were selected based on homogeneous treatment response, as assessed by expert pathologic evaluation using the College of American Pathologists (CAP) treatment effect scoring system for pancreatic cancer(39). Cases were stratified into two groups: (i) non-responders (CAP score 3), defined by minimal or no tumor response to therapy, all of whom experienced disease recurrence; and (ii) responders (CAP score 1), defined by marked tumor regression with minimal residual cancer, none of whom developed recurrence at last follow-up. Representative FFPE tissue sections from these cases were reviewed to identify blocks with the most homogeneous treatment response. Unstained sections were subsequently cut from these blocks and submitted for Visium HD spatial transcriptomics, enabling high-resolution two-dimensional evaluation of tumor regions and their associated tumor microenvironments.

### CODA 3D reconstruction of serial histology images

Scanned H&E images were downsampled to tif images at one, two, and four µm per pixel resolution for downstream processing using the Openslide platform (40). To reconstruct the microanatomical structures in human pancreatic samples into 3D volumes, CODA nonlinear image registration was employed to all image stacks at 1 µm per pixel resolution (2). Registration performance was computed by measuring tissue warp and pixel correlation between adjacent sections, with <20% tissue warp and >85% per-image pair pixel cross correlation deemed acceptable. InterpolAI was used to generate missing images to restore microanatomical connectivity (41).

To label the microanatomical components of pancreatic samples, we trained a DeepLabv3+ semantic segmentation model using the CODA workflow to recognize PDAC, non-neoplastic ducts, acini, islets, smooth muscle, endothelium, nerves, immune cell aggregates, stroma, fat, and background at 1 µm per pixel resolution (2,42). For training, a minimum of four images per 3D sample were extracted. In each image, ten to fifteen examples of each tissue type were manually annotated using Aperio ImageScope. Annotations were imported and formatted into training tiles as described in our previous method (43). Training tiles were used to finetune a deepLabv3+ semantic segmentation algorithm. One annotated image per sample was held back for model testing, with the remaining images used for model training. Following training, additional annotations were added to improve model performance until a minimum overall accuracy of 90% and a minimum per-class precision and recall of 85% was achieved. Next, the full cohort of 1 µm per pixel TIF images was segmented with this trained model. Segmented image masks were aligned by applying the transformation matrices calculated on the low-resolution H&E images to generate 3D, labelled image stacks. Volumetric matrices were generated for 3D renderings and downstream calculations and were generated at a resolution of 4 x 4 x 12 µm^3^.

StarDist instance segmentation was used to extract 20 morphological features per nucleus in the H&E images (3,4). Extracted features consisted of nuclear area, perimeter, circularity, aspect ratio, compactness, eccentricity, extent, form factor, maximum radius, mean radius, median radius, major axis length, minor axis length, orientation in degrees, red mean intensity, green mean intensity, blue mean intensity, red standard deviation in intensity, green standard deviation in intensity, and blue standard deviation in intensity. To integrate these nuclear masks with the 3D models, cell centroids were aligned using the transformation matrices calculated on the low-resolution H&E images using our previously reported method (4).

### Labelling sensitive and persistent cancer cells in 3D

Within the 3D reconstructed tumor maps containing microanatomical labels from CODA DeepLabv3+ semantic segmentation(42) and nuclear masks from StarDist instance segmentation(44), we further annotated sensitive and persistent cancer cells based on 3D connectivity. Within each volumetric matrix, we isolated connected voxels predicted to be pancreatic cancer by the segmentation model and counted the volume of each cancer cluster. We defined 0.01 mm^3^ as a threshold and defined sensitive tumor cells as all cancer clusters less than this threshold and defined persistent tumor as all cancer clusters greater than or equal to this threshold.

### Automated, 3D-guided selection of regions of interest for multi-omics analysis

We developed an algorithm to identify regions of interest (ROIs) containing an optimal mix of sensitive and persistent cancer cells where we desired to perform multi-omics profiling. The algorithm screened each section of each 3D sample within the generated volumetric matrices by extracting windows of size 6.5 x 6.5 mm^2^ with a stride of 0.8 mm. For each window we calculated the percent tissue composition of stroma and cancer within the window, as well as the percentage of the tumor that is sensitive or persistent. We kept a list of all windows containing >20% stroma (normalized by tissue area), >5% tumor (normalized by tissue area), >20% sensitive tumor (normalized by tumor area), and >20% persistent tumor (normalized by tumor area). From this list, the algorithm saved histology zoom-ins of the five windows containing the highest overall tumor percentage. These five candidate windows were reviewed by pathologists for the final selection of the optimal ROI.

Within each 6.5 x 6.5 mm^2^ ROI, we further developed an algorithm to identify four 1.5 x 1.5 mm^2^ windows containing close-to-pure compositions of either sensitive or persistent cancer cells. The algorithm independently calculated the sensitive and persistent cell density within square 1.5 x 1.5 mm^2^ windows continuously across the 6.5 x 6.5 mm^2^ ROI. Using these cell density heatmaps, the algorithm identified two non-overlapping 1.5 x 1.5 mm^2^ windows containing the highest density of sensitive cancer cells, then identified two non-overlapping 1.5 x 1.5 mm^2^ windows containing the highest density of persistent cancer cells. The larger 6.5 x 6.5 mm^2^ ROI and the four smaller 1.5 x 1.5 mm^2^ ROIs were overlaid on the corresponding H&E image to visually guide the spatial transcriptomics and spatial proteomics assays on adjacent, unstained sections.

### PIVOT co-registration and deconvolution of histology-derived features into spatial transcriptomics and spatial proteomics

We used PIVOT, a software for co-registration and integration of multi-omics datasets, to co-register and integrate the serial H&E images with the spatial transcriptomics and spatial proteomics data (45). First, PIVOT was used to register the Visium “tissue_high_res” image, the seqIF image, and each of the four IMC images to overlay on the nearest H&E image used for 3D reconstruction. Registration accuracy was deemed acceptable when a fiducial point-based root mean squared error (RMSE) less than 5 pixels was obtained per image pair, and when the registered image pairs passed visual inspection.

Following registration, the boundaries of each IMC window were overlaid on the DeepLabv3+ semantic segmentation results on the H&E image. From this, we calculated the H&E-derived tissue composition and ratio of sensitive and persistent cancer cells within each of the four IMC windows. Additionally, the Visium spots were overlaid on the DeepLabv3+ semantic segmentation and StarDist nuclear segmentation results on the H&E image. For Visium, a high-resolution scan of the same section where Visium was performed was used as the fixed image, such that the spot composition could be determined with high accuracy. We radially dilated each nuclear mask by 2 µm to estimate the cellular boundaries on H&E and determined the cell type by comparing the nuclear mask with the DeepLab segmentation results. We used this data to extract cell type and number on a per-spot basis using our previously reported method (46). Per spot cell type count and cell type composition were incorporated into the “tissue_positions.csv” file and directly imported into Seurat to define cell type labels and enable differential gene expression analysis of the histologically defined sensitive and persistent cancer cells.

### IMC data generation and analysis

Slides were stained for IMC as previously described(17). Specific tissue sections were selected based on CODA 3D models of surgically resected tumor blocks. The selected slides were baked in a pre-heated oven for 2 hours at 60°C then deparaffinized using xylene for 20 minutes. The tissues were rehydrated with an alcohol gradient (100%, 95%, 80%, and 70% EtOH for 5 minutes each). Maxpar water (Standard BioTools) was used to wash the slides before being placed in a target retrieval solution (antigen retrieval agent, pH 9, 1:10 in Maxpar water) at a sub-boiling temperature (90°C-95°C) for 1 hour. This was followed by another wash with Maxpar PBS for 10 minutes with gentle agitation. Slides were subsequently blocked using 3% BSA for 45 minutes at RT and then incubated overnight in the IMC antibody cocktail solution (**Table S2**) at 4°C. The sections were stained the next day with Cell-ID Intercalator-Ir (Standard BioTools) diluted at 1:400 in Maxpar PBS for 30 minutes at RT for nuclear counterstaining. Ruthenium tetroxide 0.5% (Electron Microscopy Sciences, PN 20700-05) at 1:2,000 was used to counterstain the tissue for 3 minutes at RT. Finally, slides were washed in Maxpar water and then air dried before being loaded in the Hyperion Imaging Plus System (Standard BioTools; RRID: SCR_023195) for image acquisition at the Johns Hopkins Mass Cytometry Facility. A total of 16 regions, two “sensitive” and two “persistent” regions from each sample, was selected for data acquisition. Upon acquisition, representative images were created using MCD Viewer 1.0560.2 (Standard BioTools; RRID: SCR_023007). Single-cell data were exported by image segmentation through a software pipeline based on DeepCell (RRID: SCR_022197) and HistoCAT 1.76 (RRID: SCR_026499). Further analysis was completed using R 4.4.0. PhenoGraph (RRID: SCR_016919) generated 43 metaclusters in an unsupervised manner which were then annotated into 7 broad cell types. Subsequently, cancer, myeloid, T, and stromal cells were further subclustered into 14, 22, 19, and 19 metaclusters that were annotated into 8, 6, 6, and 4 subtypes, respectively. Spatial influences, which incorporate both proximity and density of nearby cells, for every cell were computed using FunCN(17). Plots were generated using ggplot and igraph.

### seqIF data generation and analysis

Prior to seqIF staining on the COMET (Lunaphore) platform, formalin-fixed paraffin-embedded (FFPE) slides were preprocessed for autofluorescence clean-up as previously described (47,48). Briefly, FFPE sections were baked at 60 °C for 1 hour, deparaffinized with three xylene washes, rehydrated through a graded ethanol series, and rinsed in distilled water following standard FFPE immunofluorescence procedures. After rehydration, slides were incubated in 3% hydrogen peroxide for 10 minutes at room temperature to reduce endogenous autofluorescence, then held in PBS. Slides were subsequently rinsed and stored in PBS until antigen retrieval.

Antigen retrieval was performed using EZ-Antigen Retriever Solution 2 at 105 °C for 15 minutes in the EZ-Retriever system (XBioGenex), following the manufacturer’s recommended protocol.

After retrieval, slides were rinsed and stored in Multistaining Buffer (BU06, Lunaphore) until use. A 12-plex seqIF protocol was generated using COMET Control Software, and all reagents were loaded according to the fully automated workflow (49,50).

Nuclear counterstaining was performed using DAPI (Thermo Scientific, cat. 62248; 1:1000), applied by either a 2-minute dynamic incubation or supplementation into secondary antibody cocktails. For all staining cycles, dynamic incubation times were set to 4 minutes for primary antibody mixes and 2 minutes for secondary antibody/DAPI cocktails. All primary antibody cocktails were diluted in Multistaining Buffer (BU06, Lunaphore). Exposure times for each imaging cycle were: DAPI, 80 ms; TRITC, 400 ms; Cy5, 200 ms.

Elution was performed for 2 minutes per cycle using Elution Buffer (BU07-L, Lunaphore), followed by a 30-second quenching step with Quenching Buffer (BU08-L, Lunaphore). Imaging was carried out in Imaging Buffer (BU09, Lunaphore). Secondary antibodies included Alexa Fluor Plus 647 goat anti-rabbit (Thermo Scientific, cat. A32733; 1:200) and Alexa Fluor Plus 555 goat anti-mouse (Thermo Scientific, cat. A32727; 1:100). Upon completion of the run, the COMET Control Software generated a raw OME-TIFF file for downstream analysis.

Quality control and image visualization were conducted using the HORIZON™ Viewer (Lunaphore), including assessment of marker localization (nuclear, cytoplasmic, or membrane) and extraction image review. Post-processing of OME-TIFF images, including tissue and cell segmentation, phenotyping, and feature extraction was performed using the standardized modules of Visiopharm® as previously described (51). Mean expression values for each marker were calculated after background subtraction, and the resulting values were exported as CSV files for downstream analysis.

Cell type phenotyping was performed using the Scanpy (v1.11.3)(52) library in the Python (v3.12.10) platform following the supervised phenotyping steps outlined in lunaphore-public/**downstream-analysis-toolbox** (50)with some modifications. Briefly, the prior exported CSV files were imported into the Python environment for downstream analysis. First the unsupervised clustering of the expression table was performed to remove any cells with signals close to background such as erythrocytes. Background subtracted data for each marker was normalized by subtracting the median and dividing by the standard deviation. Dimensionality reduction was then performed using uniform manifold approximation and projection (UMAP)(53) algorithm implemented in the Scanpy framework. For UMAP n_neighbors parameter was set to 40 with a min_dist parameter set to 0.5. The Leiden clustering was performed using a resolution parameter set to 0.2. Clusters with low expression levels in all channels were considered as clusters mainly composed of cells with low expressions or cell types that was not related to any of the tested markers.

Supervised cell type assignment was performed following the steps provided in lunaphore-public/**downstream-analysis-toolbox**. The binary tree classifier shown in **Fig. S9** was used to identify the cell types. The same procedure was used to classify the tumor cells, where tumor cells expressing KERATIN 17 were identified as Basal, cells positive for GATA6 or TFF1 only were classified as Classical and cells positive for both Classical and Basal markers were classified as co-expressers.

### High-definition spatial transcriptomics

High-definition spatial transcriptomics profiling was performed using the FFPE Visium HD platform (10x Genomics), following manufacturer’s instructions, on FFPE PDAC specimens. Briefly, 5μm sections from each FFPE sample were mounted on charged frosted microscope slides and stained with H&E to facilitate the selection of regions of interest (ROIs) with tumor cells. The selection of samples and ROIs was performed by experienced pathologists (LDW and HM). The H&E-stained sections underwent decrosslinking, destaining, and probe hybridization targeting ∼18,000 human transcripts. Using the CytAssist equipment, the hybridized probes were transferred to the Visium HD slide, where they were captured, reverse transcribed into cDNA, amplified, and indexed for sequencing. Sequencing was performed at the Johns Hopkins Experimental and Computational Genomics Core (ECGC) using the NovaSeq X (Illumina) platform, with a depth of ∼350 million reads per sample following parameters recommended by 10x Genomics. This assay provided whole transcriptome profiling with single-cell resolution (2μm x 2μm).

Sequencing data preprocessing was performed using Space Ranger v3.0.1 (10x Genomics), with alignment to the human GRCh38 reference genome. The preprocessed data served as input for cell type annotations with RCTD (18) using a single-cell RNA sequencing PDAC atlas as a reference (19). The RCTD annotated data was subsequently analyzed using the R/Bioconductor package Seurat (54).

PDAC subtypes classification was performed using previously established signatures for the classical and basal-like subtypes (55) using the Seurat *AddModuleScore* function. PDAC spots were classified as classical if the score was 20% higher than the basal score, and vice-versa. Spots whose scores did not fit the criteria were classified as intermediate.

Differential expression analysis and gene set enrichment analysis for MSigDB Hallmark Pathways were performed on the PDAC pseudo-bulked data to compare transcriptional changes between non-responding and responding tumors.

### Ligand-receptor interaction analysis using SpaceMarkers

Ligand-receptor (L-R) interactions from CAFs to PDAC cells were quantified using SpaceMarkers version 1.99.0 (56) applied to the Visium HD data to identify interactions associated with lack of chemotherapy response. The analysis utilized the CellChatDB.human ligand-receptor interactions database from CellChat v2.1(57). With the spatial distributions of CAFs and PDAC determined through RCTD, the influence of both cell types was modeled using spatial kernel-based smoothing with the *calculate_influence* function, with the kernel width set to 16 μm. Next, we applied the *calculate_thresholds* function in SpaceMarkers to identify thresholds for determining hotspots of PDAC presence and CAF influence via two-component Gaussian mixture models based on spatial distributions and influences of both cell types. Hotspots of CAF presence and PDAC influence were subsequently annotated using the *find_hotspots* function, where any bin exhibiting an influence hotspot for a given cell type was designated as being under that cell type’s influence.

We further stratified hotspots of CAF presence by the influence of PDAC and performed a t-test to identify differentially expressed genes in CAF hotspots influenced by PDAC compared to CAF hotspots without PDAC influence. Scores for each gene were computed using the *calculate_gene_scores_directed* function employing Cohen’s D effect size from the effsize package v0.8.1. Ligand complex scores in CAFs were obtained by performing a weighted average of the scores of associated genes, weighted by their frequency of occurrence in the data utilizing the *calculate_gene_set_score* function. Receptor complex scores for PDAC were calculated using the *calculate_gene_set_specificity* function, assigning a specificity score to each gene in the receptor complex based on the log-fold change between the bins in the top and bottom quintiles of PDAC presence. The ligand-receptor interaction score was determined as a weighted geometric mean of the ligand and receptor scores through the *calculate_lr_scores* function. After stratifying samples by response, we calculated the effect size of the L-R score change from responders to non-responders, ranked the L-R pairs by decreasing effect size, and plotted a heatmap of the top 30 L-R pair scores.

### Standard spatial transcriptomics

For the Visium standard (Visium SD) approach, using Visium CytAssist FFPE platform (10x Genomics) following the manufacturer’s protocol, ROIs were selected based on cellular composition as determined during CODA 3D reconstruction. The 5μm sections of each sample underwent deparaffinization and H&E staining. Human probe hybridization was performed overnight at 50°C, followed by cDNA synthesis, amplification, and indexing. The probe set was identical to that used for the Visium HD profiling. All libraries were sequenced to a depth of ∼250 million reads per sample on the NovaSeq X (Illumina) at the ECGC. The Visium SD approach provided whole transcriptome profiling with a resolution of 55μm.

Sequencing data preprocessing was performed with Space Ranger v2.0.1 using GRCh38 as the reference genome. Downstream analysis was performed in Seurat, with cell types captured in each Visium SD spot classified by CODA. For each 55μm spot, CODA quantified the detected cell types according to their morphological characteristics. In Seurat, the final classification for each spot was determined by the cell type that represented at least 51% of the spot. Spots in which none of the detected cell types achieved this threshold were classified as mixed. PDAC subtype classification used the same strategy as described for the Visium HD dataset. Following cell type classification, we performed downstream analysis to validate the ligand-receptor interactions identified by SpaceMarkers. Initially, we identified PDAC spots that exhibited increased expression of CAF markers to select spots with higher probability of capturing CAF-PDAC interactions. Subsequently, we verified the expression of CAF ligands and PDAC receptors identified in the Visium HD analysis as drivers of stroma-to-tumor signaling.

## Author contributions

LW, AK, WJH, LK, EF, and DW conceived the study. AF led the 3D reconstruction and multi-omics integration. HM and BAP led the cohort design and clinical correlates. MW, DL, and AD led the spatial transcriptomics analysis. AV, SMS, SP, KIR, and PAG led the spatial proteomics analysis. All authors read and approved the manuscript.

## Data and code availability

The CODA software for 3D reconstruction of serial histological images is available at the following link: https://github.com/ashleylk/CODA. The PIVOT software for co-registration of spatial proteomics and transcriptomics with histology is available at the following link: https://github.com/Kiemen-Lab/CODA_pivot. The software for IMC analysis is available at the following link: https://github.com/wjhlab/BTC_chemo and https://github.com/wjhlab/CAF-neighborhoods-PDAC. The software for spatial transcriptomics analysis is available at the following link: https://github.com/FertigLab/dpt-paper-2026. The software for seqIF analysis is available at the following link: https://github.com/lunaphore-public/downstream-analysis-toolbox.

## Acknowledgments

The authors acknowledge the following sources of funding:

Break Through Cancer; American Association of Cancer Research; Lustgarten Foundation; NIH/NCI U54CA268083; NIH/NCI U24CA284156; NIH/NCI T32CA193145; NIH/OD S10OD034407

---

**Figure S1.**
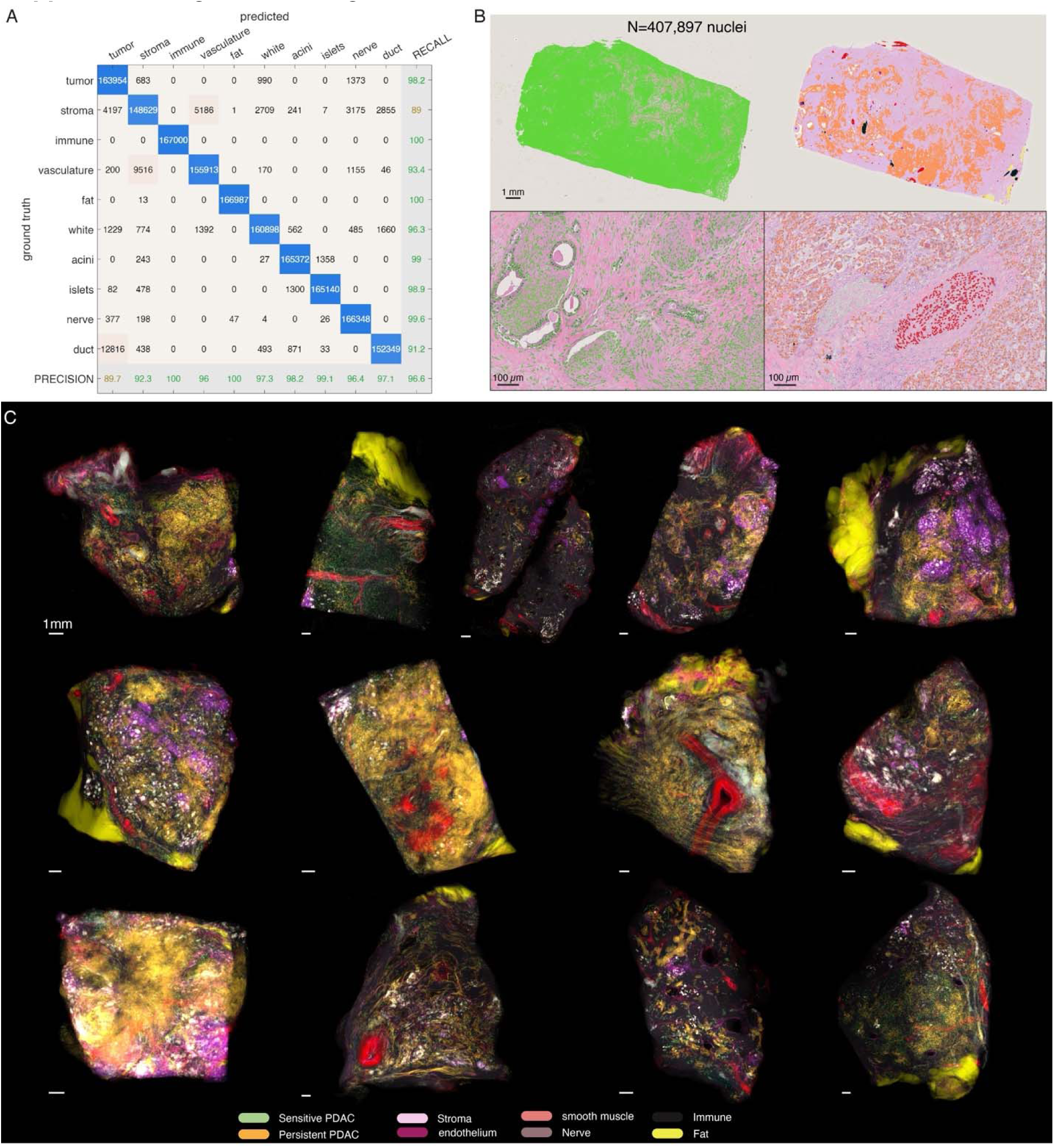
Validation of semantic segmentation model and nuclear feature extraction pipeline. **(a)** Performance metrics for the semantic segmentation model trained to recognize ten pancreatic microanatomical components, achieving 96.6% overall accuracy in independent testing images. **(b)** StarDist-based nuclear segmentation and extraction of 20 morphological features per nucleus, including area, major axis length, and color. **(c)** Z-projections of the 3D cellular-resolution maps show a high degree of heterogeneity within each sample.

**Figure S2.**
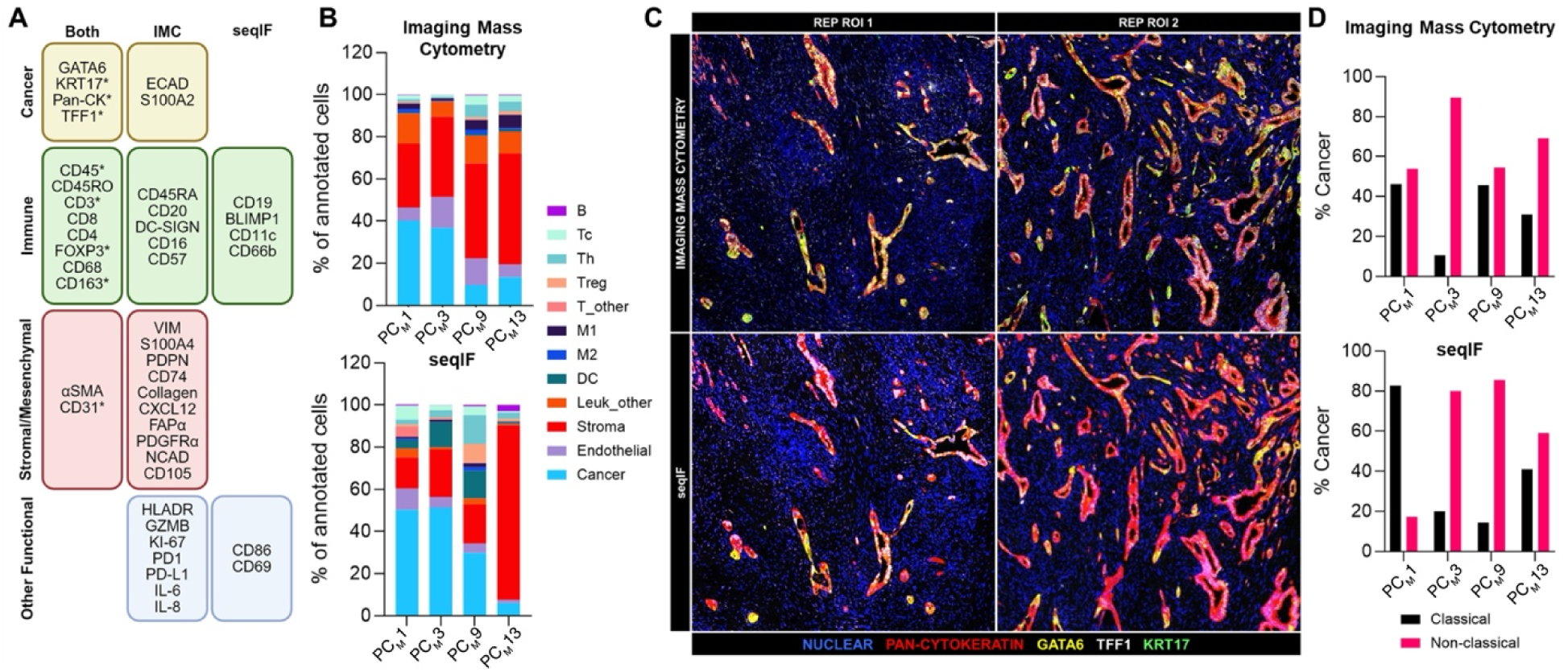
Cross validation of high-parameter single-cell protein-level analysis. **(a)** Comparison of antibody panels used for IMC and seqIF. *Denotes different antibody clones. **(b)** Global cell type compositional profiles delineated by IMC and seqIF. **(c)** Representative images for cancer cell subtyping markers detected by IMC and seqIF from two different regions of interest (ROI). Scale bar: 200 µm. **(d)** Proportions of classical or non-classical (mixed, basal, and subtypes not otherwise specified) cancer subtypes delineated by IMC and seqIF are shown for each tumor specimen. Abbreviations: Tc, CD8^+^ T cells; Th, CD4^+^ helper T cells, Treg; FOXP3^+^ regulatory T cells, T_other; CD3^+^CD4^-^CD8^-^ T cells; M1 and M2, macrophage clusters; DC, dendritic cells; Leuk_other, other leukocyte clusters.

**Figure S3.**
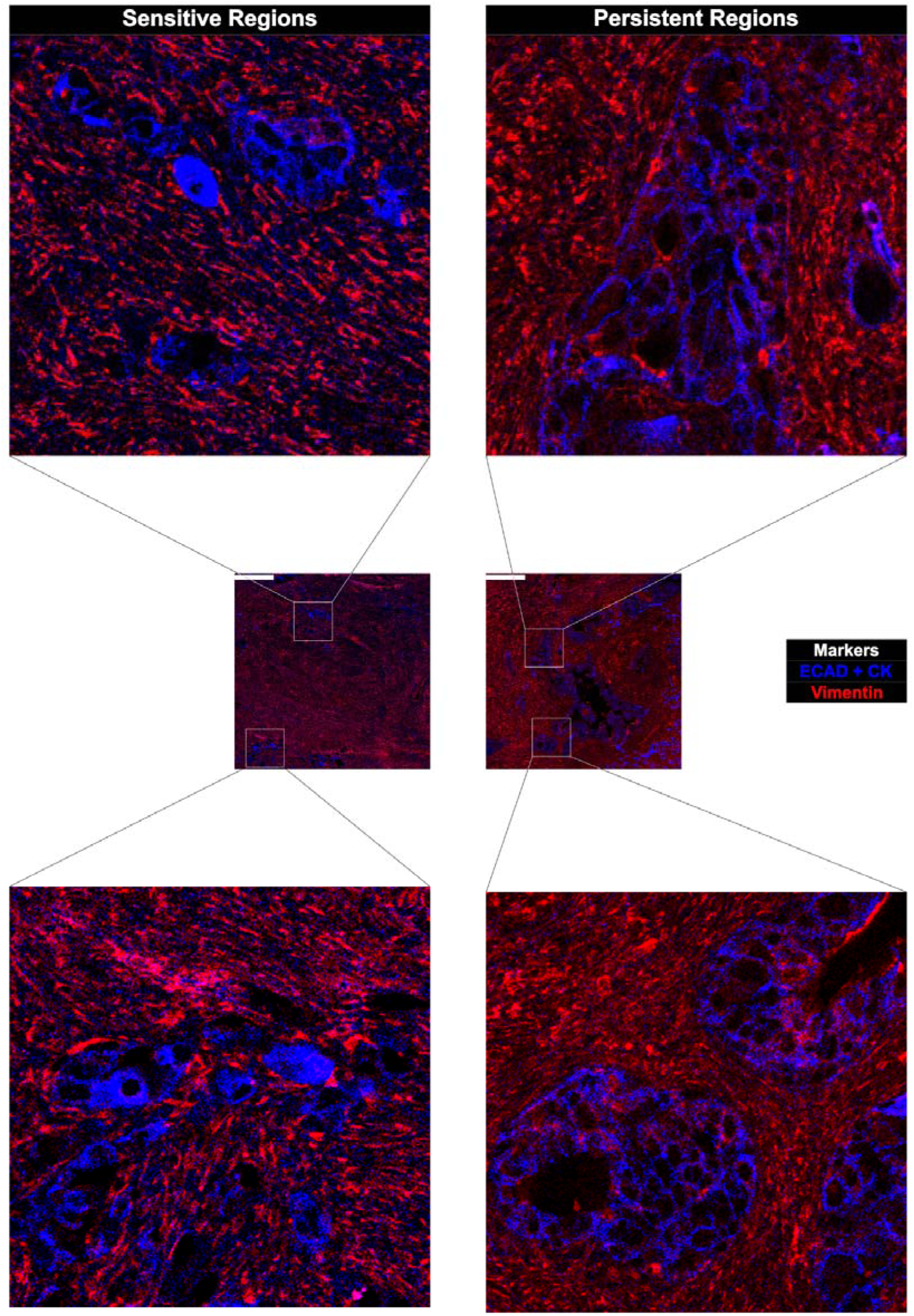
Representative IMC images for high vimentin expression by sensitive and persistent cancer cells. **Scale bar: 300 µm.**

**Figure S4.**
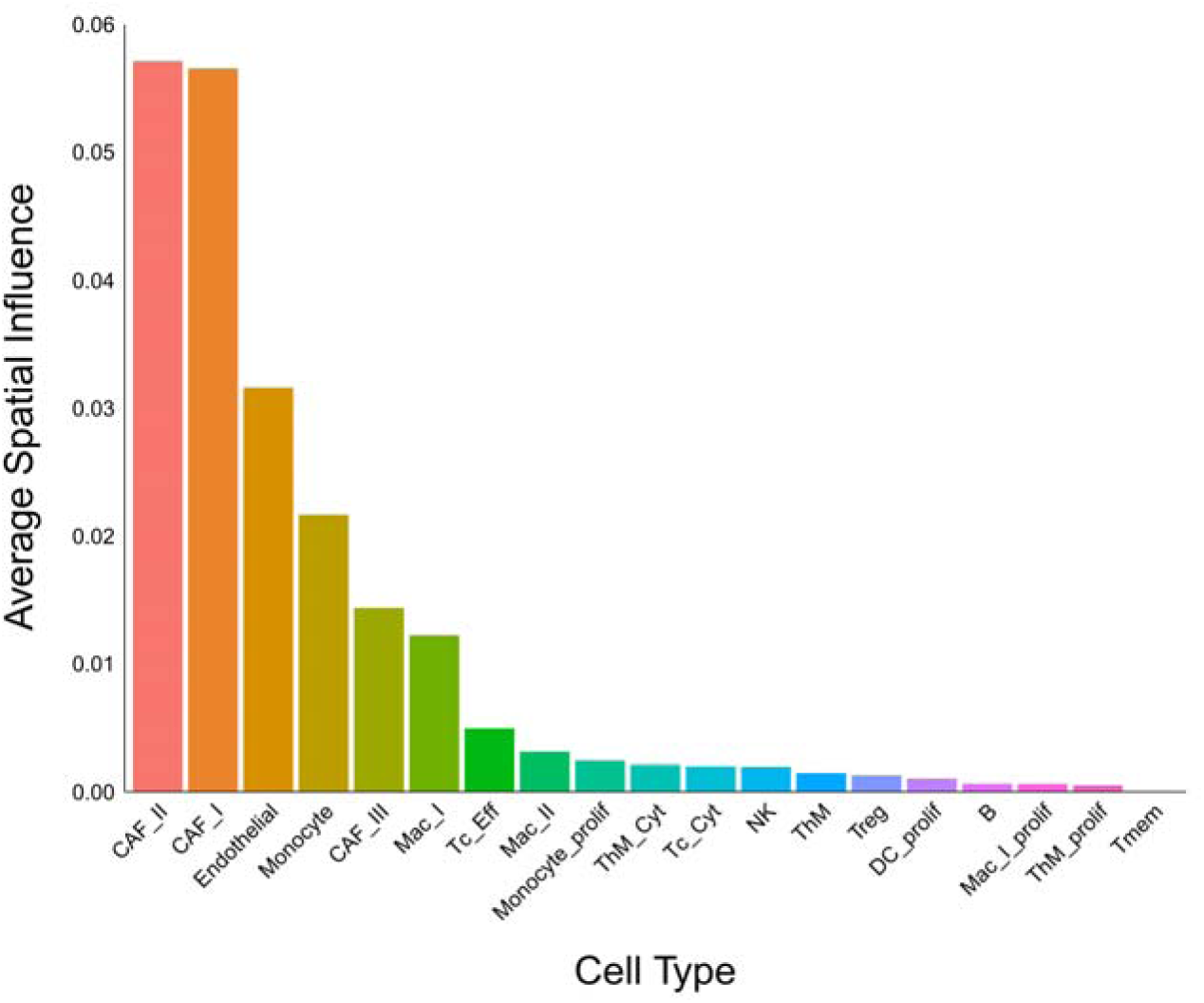
Average spatial influence on cancer cells from nearby cell types. Spatial influence of 1 indicates the totality of spatial influences exerted onto a given cell by neighboring cells based on their proximity and frequency. Abbreviations: Tc, cytotoxic CD8^+^ T cells; Th, helper CD4^+^ T cells; Tc_Cyt, GZMB^int^; Tc_Eff, CD45RA^+^GZMB^hi^; ThM, CD4^+^CD45RO^+^ memory T cells; Tmem, CD3^+^CD4^-^CD8^-^ memory T cells; Treg, CD4^+^FOXP3^+^ regulatory T cells; NK, CD57^+^ natural killer; DC, dendritic cells; Mac, macrophages; CAF, cancer-associated fibroblasts; prolif, Ki67^+^ proliferating; NOS, not otherwise specified; UA, unassigned.

**Figure S5.**
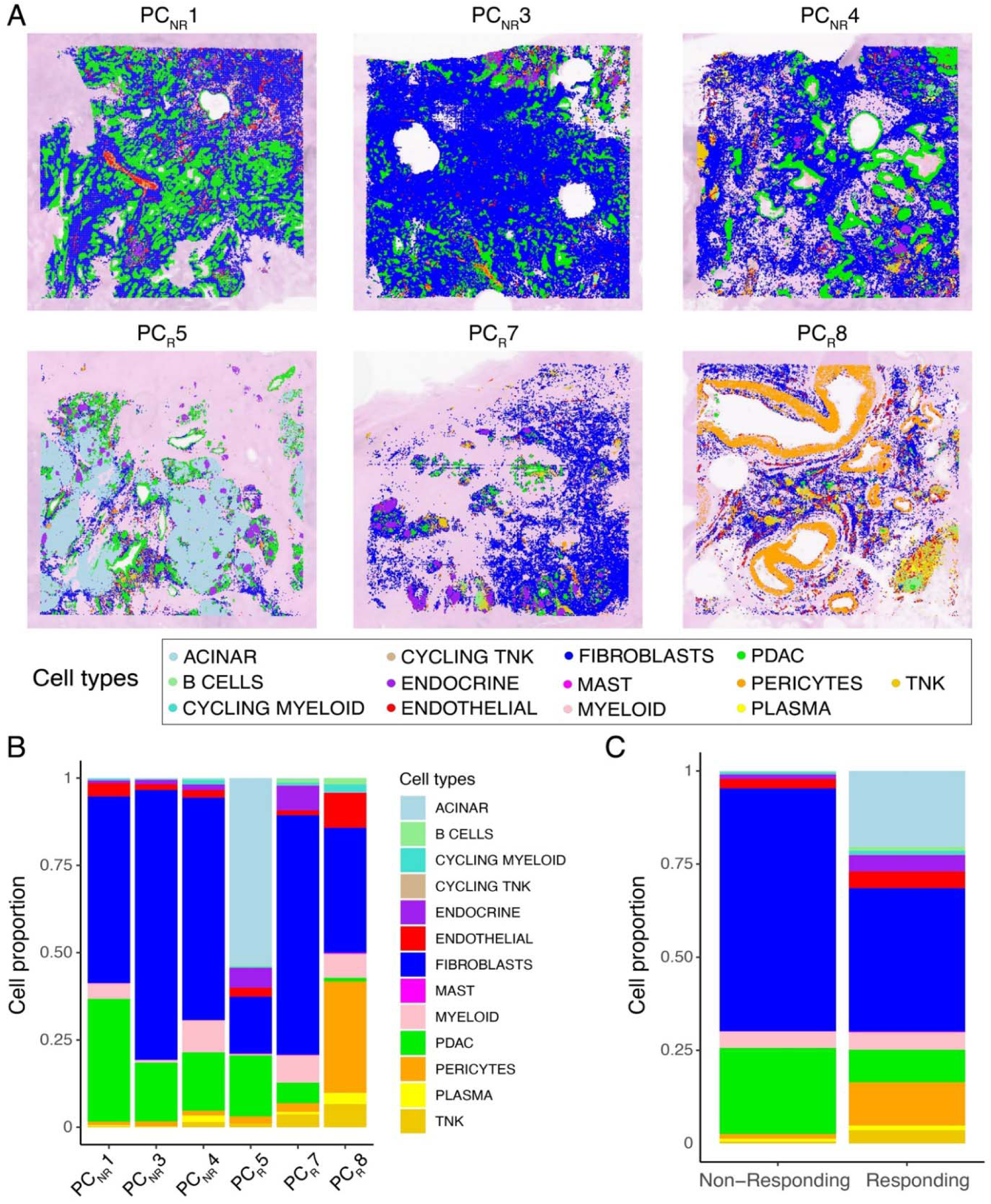
Cell types detected on Visium HD spatial transcriptomics. **(a)** Spatial distribution of cell types across all samples (NR - non-responding; R - responding). **(b)** Proportion of cell types detected in each sample. **(c)** Proportion of cell types in non-responding and responding samples.

**Figure S6.**
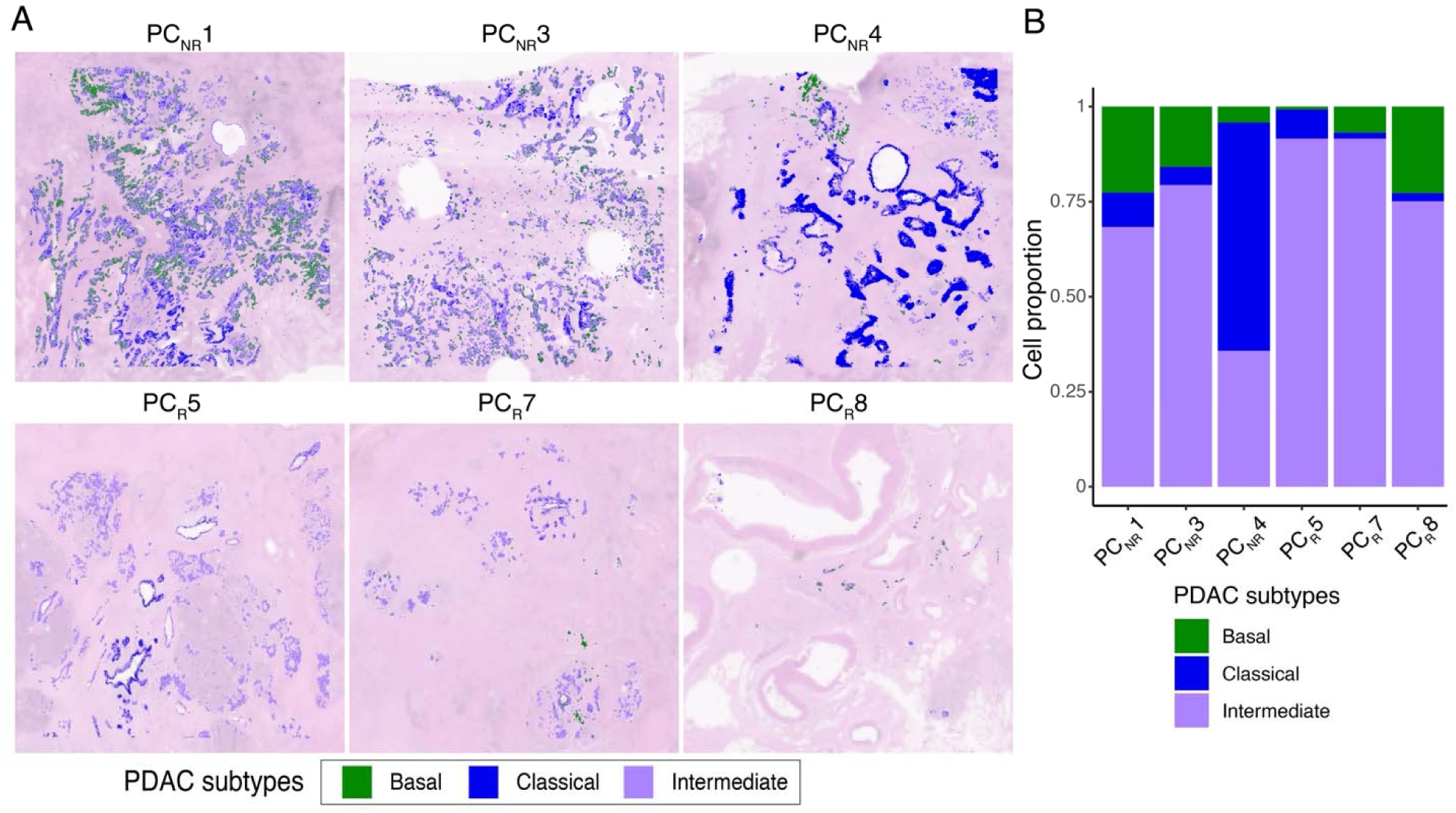
PDAC transcriptional subtypes detected Visium HD spatial transcriptomics. **(a)** Spatial distribution of PDAC subtypes across all samples (NR – non-responding; R - responding). **(b)** Proportion of PDAC subtypes detected in each sample.

**Figure S7.**
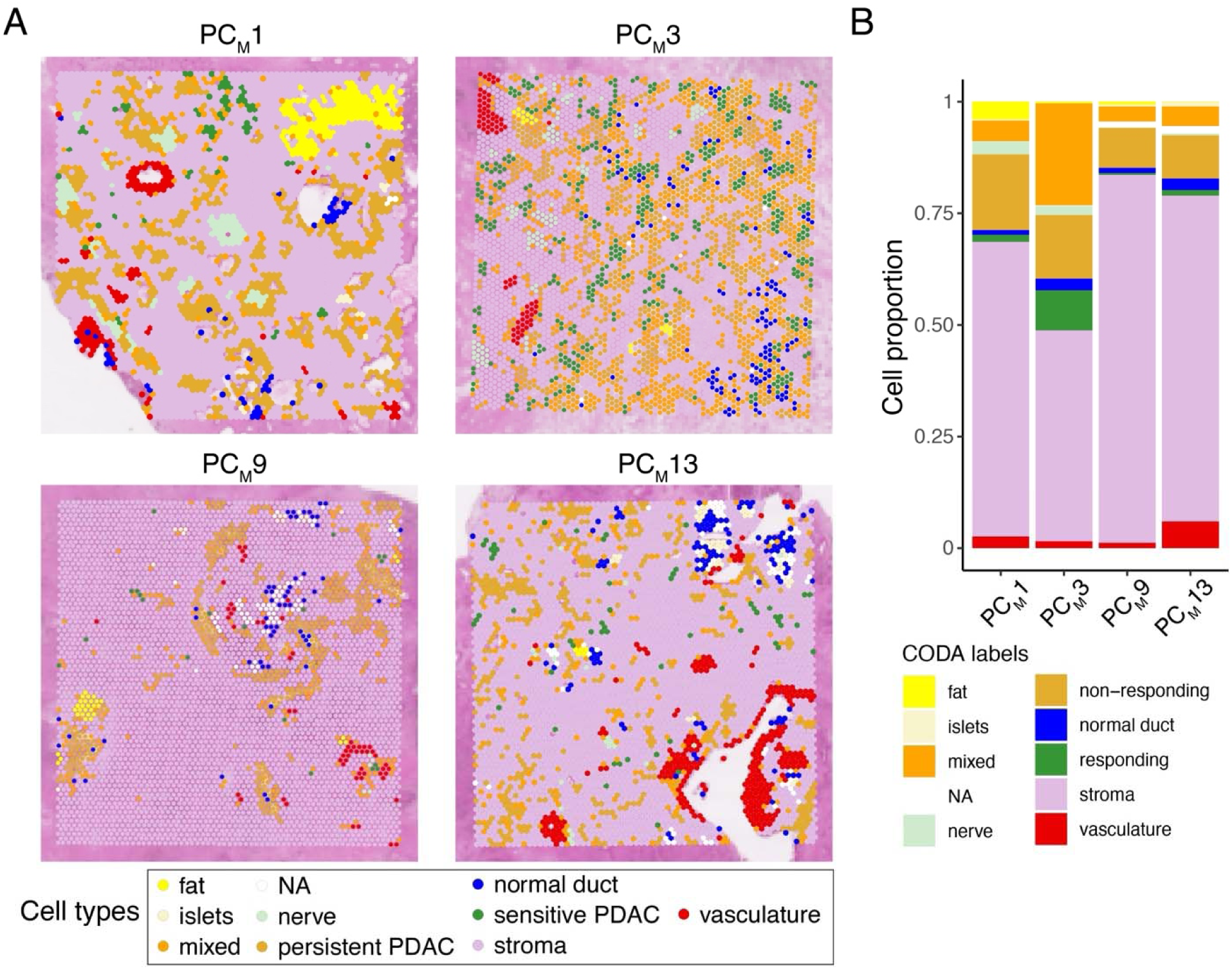
Cell types detected Visium SD spatial transcriptomics on 3D-mapped samples. **(a)** Spatial distribution of cell types across all samples. **(b)** Proportion of cell types detected in each sample.

**Figure S8.**
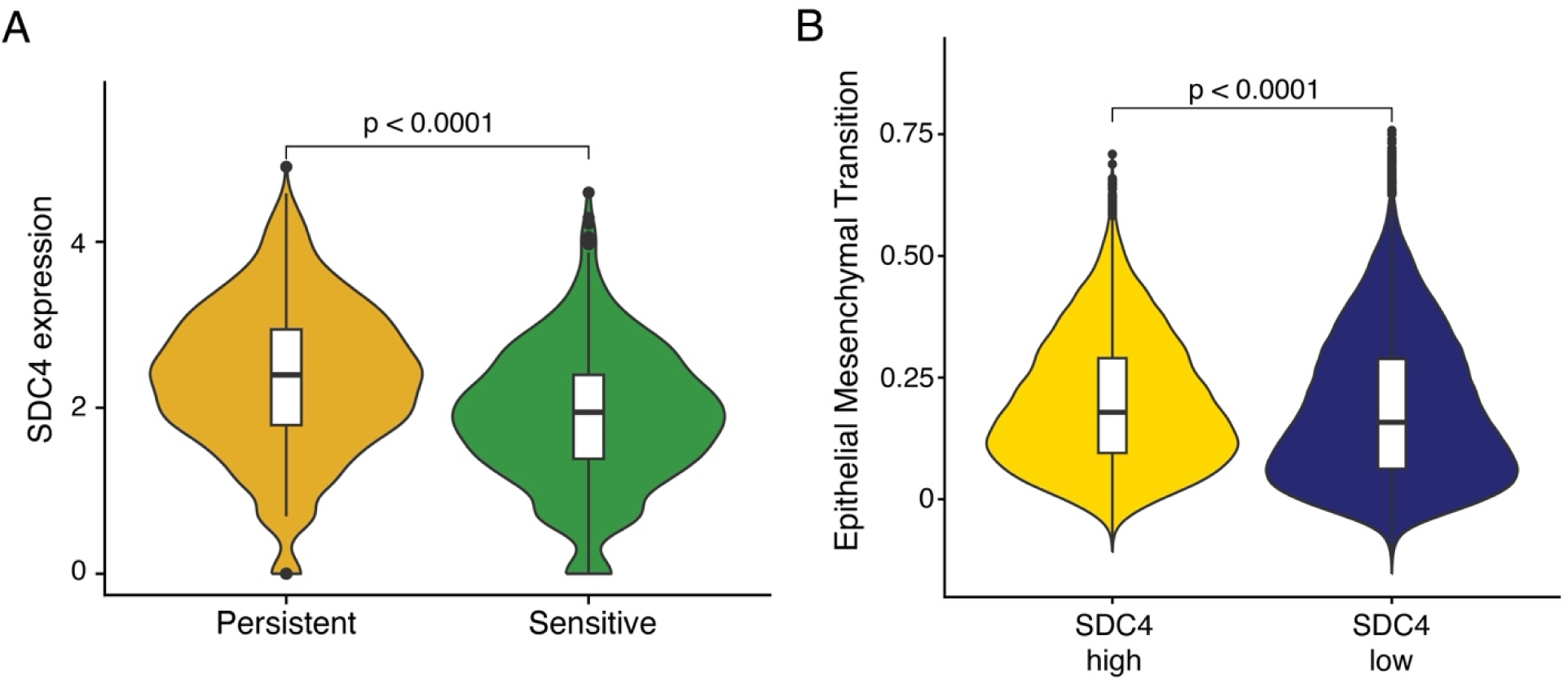
SDC4 expression on Visium SD CAF and PDAC mixed spots in 3D-mapped samples. **(a)** SDC4 expression in CAF-PDAC mixed spots in persistent and sensitive tumor spots. **(b)** Expression of EMT pathways in SDC4 high and low samples.

**Figure S9.**
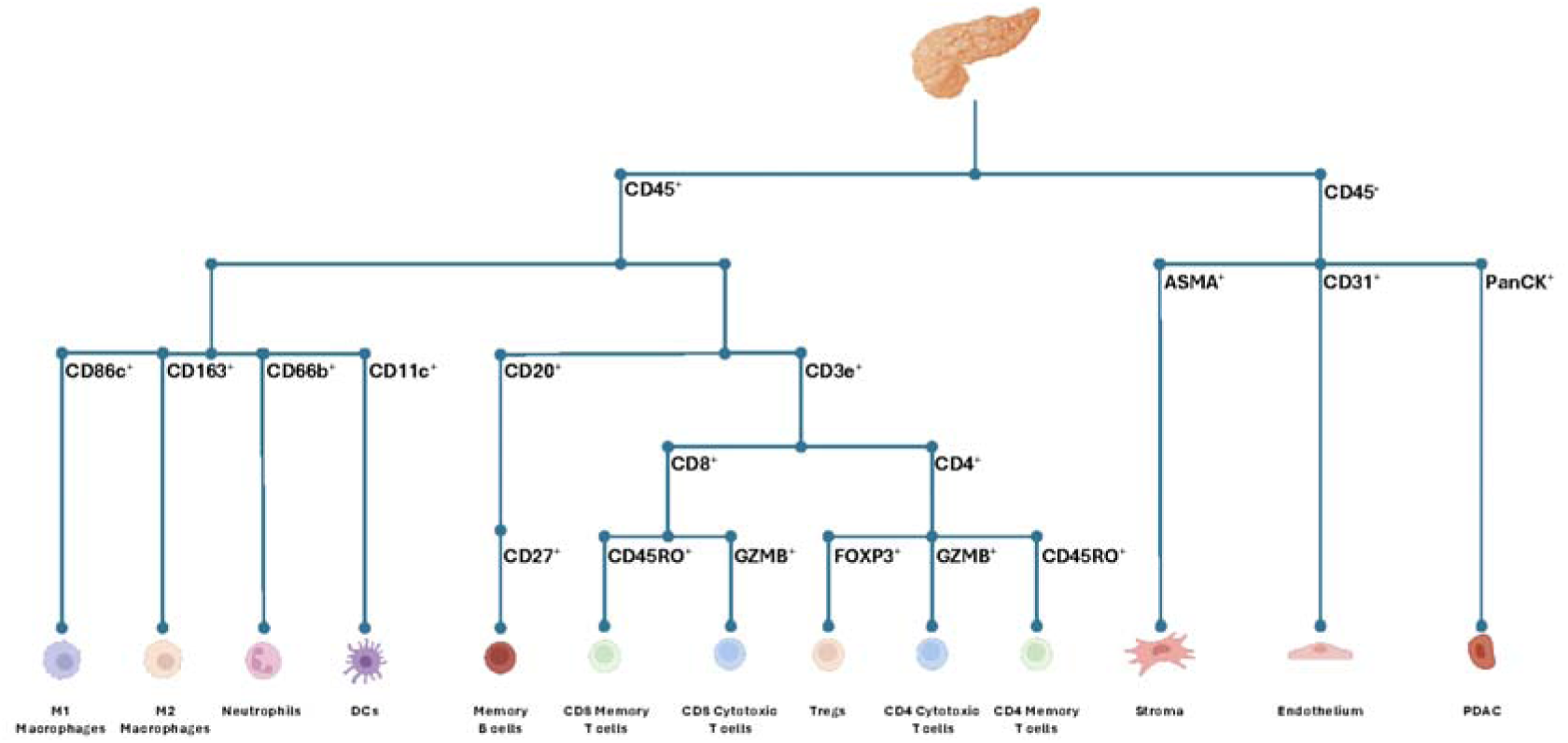
**Cell type classification tree for seqIF analysis.**

**Table S1.** Patient cohort data.

| Samples | Age (yrs) | sex | TNM stage | Tumor Grade | Neoadjuvant treatment | Neoadjuvant Treatment Regimen |  |  | Treatment effect | Recurrence status | Recurrence timing (days after initial resection) | Follow-up length (days after initial resection) |
| --- | --- | --- | --- | --- | --- | --- | --- | --- | --- | --- | --- | --- |
|  |  |  |  |  |  | FOLFIRI-NOX | Gemcitabine/nab-paclitaxel | Other |  |  |  |  |
| PC<sub>M</sub>1 | 70s | F | ypT2 pN0 | G3 | yes | yes | yes | yes | Score 2 | No |  | 91 |
| PC<sub>M</sub>3 | 70s | F | ypT2 pN2 | G2 | yes | yes | yes | no | Score 3 | Yes | 918 |  |
| PC<sub>M</sub>5 | 80s | M | ypT2 pN2 | G2 | yes | yes | yes | no | Score 2 | No |  | 120 |
| PC<sub>M</sub>9 | 60s | M | ypT3 pN1 | G3 | yes | yes | no | no | Score 2 | Yes | 228 |  |
| PC<sub>M</sub>10 | 60s | M | ypT3 pN1 | G2 | yes | yes | yes | yes | Score 2 | Yes | 271 |  |
| PC<sub>M</sub>11 | 70s | F | ypT3 pN1 | G3 | yes | no | yes | yes | Score 3 | Yes | 437 |  |
| PC<sub>M</sub>13 | 50s | M | ypT2 pN1 | G3 | yes | yes | no | no | Score 2 | Yes | 582 |  |
| PC<sub>U</sub>14 | 50s | M | pT2 pN1 | G3 | no | no | no | no | N/A | Yes | 237 |  |
| PC<sub>U</sub>15 | 60s | F | pT2 pN0 | G2 | no | no | no | no | N/A | Yes | 316 |  |
| PC<sub>M</sub>16 | 40s | F | ypT1c pN0 | G2 | yes | yes | no | no | Score 2 | Yes | 204 |  |
| PC<sub>M</sub>19 | 50s | F | ypT3 pN0 | G3 | yes | yes | no | yes | Score 2 | No |  | 794 |
| PC<sub>M</sub>23 | 80s | M | ypT2 pN1 | G2 | yes | no | yes | no | Score 3 | Yes | 214 |  |
| PC<sub>M</sub>25 | 70s | M | ypT2 pN1 | G3 | yes | yes | no | no | Score 2 | Yes | 582 |  |
| Samples | Age (yrs) | sex | TNM stage | Tumor Grade | Neoadjuvant treatment | Neoadjuvant Treatment Regimen |  |  | Treatment effect | Recurrence status | Recurrence timing (days after initial resection) | Follow-up length (days after initial resection) |
|  |  |  |  |  |  | FOLFIRI-NOX | Gemcitabine/nab-paclitaxel | Other |  |  |  |  |
| PC<sub>NR</sub>1 | 50s | M | ypT4 pN2 | G3 | yes | yes | no | no | Score 3 | Yes | 135 |  |
| PC<sub>NR</sub>3 | 60s | F | ypT2 pN0 | G2 | yes | yes | no | no | Score 3 | Yes | 268 |  |
| PC<sub>NR</sub>4 | 80s | F | ypT2 pN0 | G1 | yes | yes | no | no | Score 3 | Yes | 344 |  |
| PC<sub>R</sub>5 | 70s | F | ypT1b pN0 | G2 | yes | yes | no | no | Score 1 | No |  | 1904 |
| PC<sub>R</sub>7 | 80s | M | ypT1 pN1 | G2 | yes | no | yes | yes | Score 1 | No |  | 167 |
| PC<sub>R</sub>8 | 60s | M | ypT1b pN1 | G2 | yes | yes | no | no | Score 1 | No |  | 1764 |

**Table S2.**
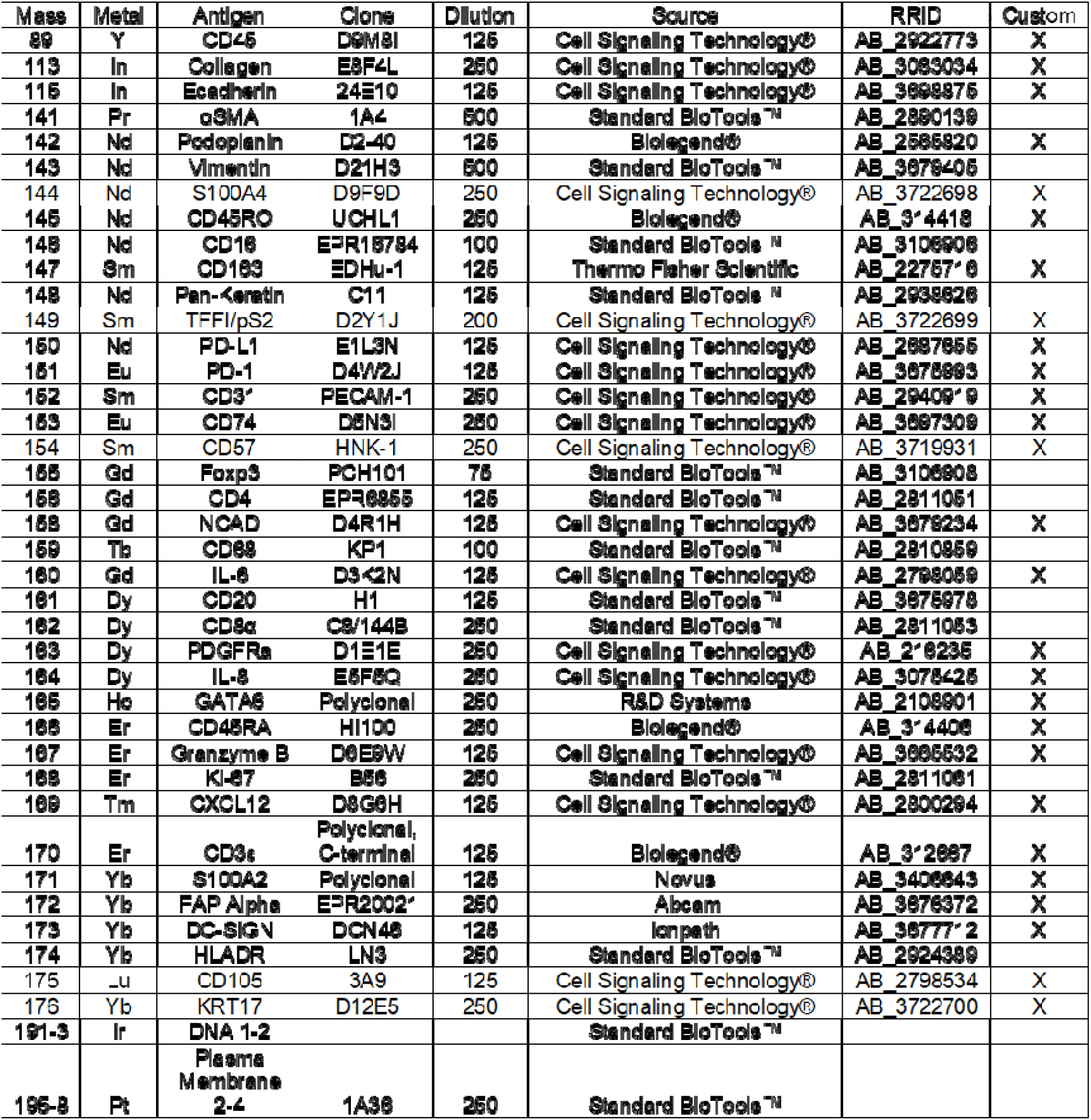
IMC antibody panel.

